# Cortical astrocyte N-Methyl-D-Aspartate receptors influence whisker barrel activity and sensory discrimination

**DOI:** 10.1101/2023.01.08.523173

**Authors:** Noushin Ahmadpour, Meher Kantroo, Michael J. Stobart, Tania Salamovska, Finnegan O’Hara, Dustin Erickson, Sofia Carrion-Falgarona, Jillian L. Stobart

**Affiliations:** College of Pharmacy, University of Manitoba, Winnipeg, MB, Canada; Centre on Aging, University of Manitoba, Winnipeg, MB, Canada

**Author notes:** Corresponding Author: Jillian Stobart, College of Pharmacy, University of Manitoba, 750 McDermot Ave. Winnipeg, MB, R3E 0T5.

## Abstract

Cortical astrocytes encode sensory information through their calcium dynamics, but it remains unclear if modulation of astrocyte calcium transients can change somatosensory circuits and behaviour *in vivo*. Here, we used a novel knockdown approach to selectively reduce astrocyte N-methyl-D-aspartate receptors (NMDAR). We found that these ionotropic receptors contribute to astrocyte Ca^2+^ transients encoding sensory information. This was essential for the optimal processing of sensory information in nearby neurons, since a reduction in astrocyte NMDARs caused circuit dysfunction and impaired neuronal responses to stimulation. This led to sensory discrimination deficits in the animal. Overall, our findings show that astrocytes can rapidly respond to glutamatergic transmission via their NMDAR and these receptors are an important component for astrocyte-neuron interactions that regulate cortical sensory discrimination *in vivo*.

## Introduction

Astrocytes are glial cells that are essential for homeostasis and have increasingly been shown to influence animal behaviour ^1–5^. Morphologically, astrocytes integrate into circuits through their highly branched processes that enwrap nearby synapses. Here, astrocytes can modulate neuronal activity through the Ca^2+^-mediated release of signaling molecules (i.e. gliotransmitters ^6,7^). Since astrocytes form an interconnected glial network and a single astrocyte contacts multiple synapses ^6,8^, local activity sensed by an astrocyte is potentially relayed to influence neurons at remote sites and at the population level ^6,9^. However, the conditions and pathways involved in the astrocytic modulation of neuronal populations and animal behaviour are not clearly understood, particularly within the cortex.

An essential component of the bidirectional crosstalk between astrocytes and neurons is astrocytic calcium signaling ^7^. Astrocyte Ca^2+^ dynamics primarily occur as Ca^2+^ microdomains (MDs), where intracellular Ca^2+^ is elevated within subcellular compartments such as the soma or fine processes^10–14^. Following nearby synaptic activity, such as that induced by somatosensory whisker stimulation, there is an increase in the number of astrocyte Ca^2+^ MDs, as well as their size (µm^2^) and signal amplitude ^10,11,13,15^. The majority of astrocyte Ca^2+^ events evoked by local circuit activity have a delayed signal onset relative to the dynamics of nearby neurons ^10,16,17^. However, several studies have reported the presence of fast-onset astrocyte Ca^2+^ MDs on the spatial scale of neuronal spines that closely parallel the time course of neuronal Ca^2+^ events ^10,15^. Therefore, astrocyte Ca^2+^ signaling has the necessary temporal and spatial properties for rapid synaptic modulation.

Numerous studies have shown that G-protein coupled receptors (Gq-GPCRs) via inositol-1,4,5-trisphosphate (IP_3_) type 2 receptors (IP_3_R2) contribute to astrocyte Ca^2+^ transients, often downstream of neuromodulators such as norepinephrine and acetylcholine ^5,10,12,18–23^. However, approaches to disrupt the Gq-GPCR pathway, such as using an inhibitory peptide (iβARK) or knockout of IP_3_R2, reduce the number of astrocyte MDs, but do not prevent astrocyte responses to whisker stimulation ^5,10,12,13^. IP_3_-mediated pathways contribute more to delayed onset astrocyte Ca^2+^ MDs than to fast onset MDs following whisker stimulation ^10^. Currently, the signaling pathways that underlie fast onset Ca^2+^ events in astrocytes remain unknown.

A growing body of evidence suggests that ion channels and ionotropic receptors can mediate astrocyte Ca^2+^ MDs via transmembrane Ca^2+^ flux ^14,24,25^. Astrocytes express ionotropic N-methyl-D-aspartate receptors (NMDARs), which are activated by synaptically released glutamate in the presence of D-serine or glycine. Studies of astrocyte NMDARs have focussed on *in vitro* preparations or brain slices^26,27,28–35,36–38^, and indicate that these receptors are primarily composed of GluN1 as the essential subunit, paired with GluN2C, 2D or GluN3 subunits ^33,37,39^ that are not sensitive to blockade by Mg^2+^ ions^36,37^. When activated, these receptors induce astrocyte membrane depolarizations, inward currents and increases in intracellular Ca^2+^ ^33,36–38,40^. NMDAR pharmacological approaches have suggested that these receptors do not mediate resting astrocyte Ca^2+^ MDs in the hippocampus ^41^, but contribute to stimulation-evoked astrocyte Ca^2+^ increases in multiple brain regions ^27,42,43^ and perivascular endfeet ^17,44^. However, all of these studies used drugs that also inhibited neuronal NMDARs, confounding the interpretation of these results.

In this study, we aimed to determine the contribution of astrocyte NMDARs to Ca^2+^ microdomains and their impact on nearby neuronal activity during whisker stimulation in awake mice *in vivo*. We implemented a novel miRNA-adapted shRNA (shRNA^mir^) strategy to knockdown (KD) the gene for the essential GluN1 NMDAR subunit, Grin1, selectively in cortical astrocytes of the barrel cortex (i.e. reduced Grin1 expression would eliminate all astrocyte NMDARs irrespective of their subunit composition). We report a novel mechanism where astrocyte NMDARs regulate Ca^2+^ MDs evoked by whisker stimulation and play a key role in regulating optimal neuronal activity needed for information processing that dictates sensory discrimination.

## Results

### Knockdown of astrocyte NMDAR expression

To reduce the expression of NMDARs in cortical astrocytes, we used an adeno-associated virus (AAV9) containing the gfaABC1D promoter, three unique miRNA-adapted shRNA hairpins for knockdown of Grin1 mRNA (Grin1 KD), and a membrane-tagged genetically-encoded calcium indicator (GECI), Lck-GCaMP6f (AAV9-sGFAP-Grin1-shRNA^mir^-Lck-GCaMP6f; Fig. 1A). A comparable AAV9 containing three non-silencing shRNA^mir^ was used as a control (AAV9-sGFAP-NS-shRNA^mir^-Lck-GCaMP6f; Fig. 1A). Similar viral constructs have been widely adopted to target various astrocyte pathways ^5,10,12,45^. Injection of these AAVs into the whisker-barrel somatosensory cortex of adult C57BL/6NCrl mice (Fig. 1B) resulted in astrocyte-specific expression of Lck-GCaMP6f (Fig. 1C, D) in more than 95% of astrocytes in the viral transduction area (Fig. 1E, F; Supplementary Fig. 1A, B), without notable astrogliosis (Stobart et al., 2018b). When we dissociated cortical tissue and collected astrocytes by flow cytometry using their GCaMP6f fluorescence (Supplementary Fig. 1C, D), we found that the expression of Grin1 was reduced at the mRNA level in the Grin1 KD astrocytes by 70% (Fig. 1G).

**Figure 1.**
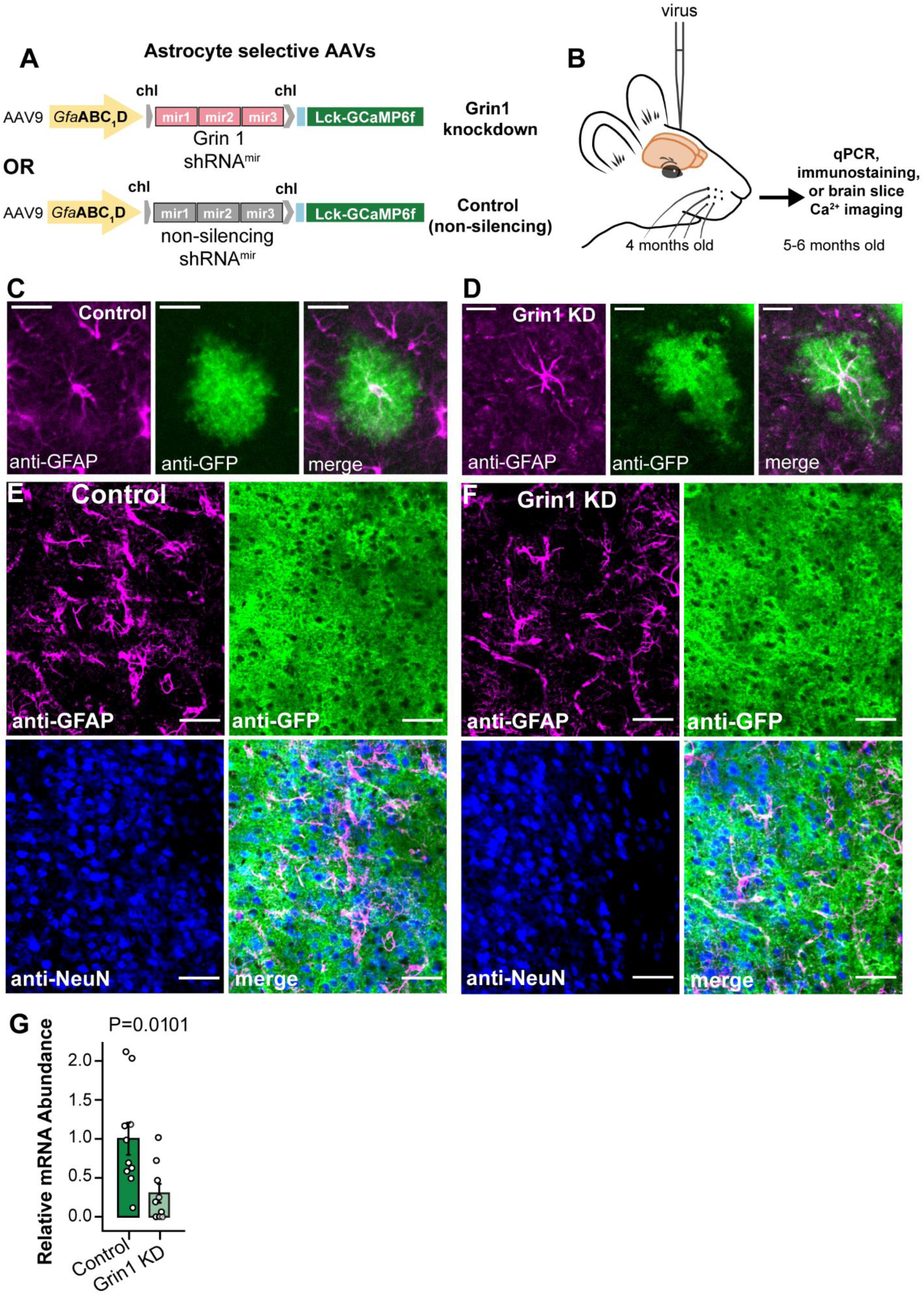
Astrocyte-specific knockdown of Grin1, the gene for the essential subunit of NMDA receptors. A) Astrocyte-specific AAV9 constructs with three shRNA^mir^ targeting Grin1 for knockdown (top) or three non-silencing shRNA^mir^ (middle) with membrane tagged Lck-GCaMP6f. B) AAVs were injected into the whisker barrel cortex. C, D) Lck-GCaMP6f (anti-GFP) from each construct localized to astrocytes (anti-GFAP) by immunohistochemistry. Scale= 25 µm E, F) Astrocytes (anti-GFAP) and not neurons (anti-NeuN) expressed the viral constructs. Scale= 50 µm G) qPCR for Grin1 mRNA abundance from isolated astrocytes showed 70% reduced expression in Grin1 KD. Mean ± SEM compared by Mann-Whitney-Wilcoxon test. n= 10 control, 9 Grin1 KD mice.

Grin1 KD also resulted in a functional reduction in astrocyte NMDAR activity evoked by agonist application to acute adult brain slices (Fig. 2A). Localized Ca^2+^ signals in active regions of interest (ROIs) were identified in Lck-GCaMP6f positive astrocytes (Fig. 2B, C) in the presence of neuronal blockers (1µM TTX, 10µM CNQX, 100µM CdCl_2_). Following bath application of NMDAR agonists, NMDA (50 µM) and D-serine (10 µM), Ca^2+^ increased in control astrocytes (Fig. 2C). This response was attenuated in Grin1 KD astrocytes (Fig. 2D), both in terms of the magnitude of the response (area under the curve; mean ± SEM; 641.65 ± 77.81 control vs. 70.68 ± 15.53 KD; Fig. 2E) and the number of Ca^2+^ peaks in each ROI (14.11 ± 3.01 control vs. 6.61 ± 2.12 KD peaks/ROI/min; Fig. 2F). Astrocytes were viable following NMDA and D-serine application because both control and Grin1 KD cells responded to alpha-1 adrenergic receptor agonist, phenylephrine, which induces IP_3_-mediated release of Ca^2+^ from intracellular stores (5.67 ± 1.14 control vs. 5.07 ± 1.17 KD peaks/ROI/min; Fig. 2G).

**Figure 2.**
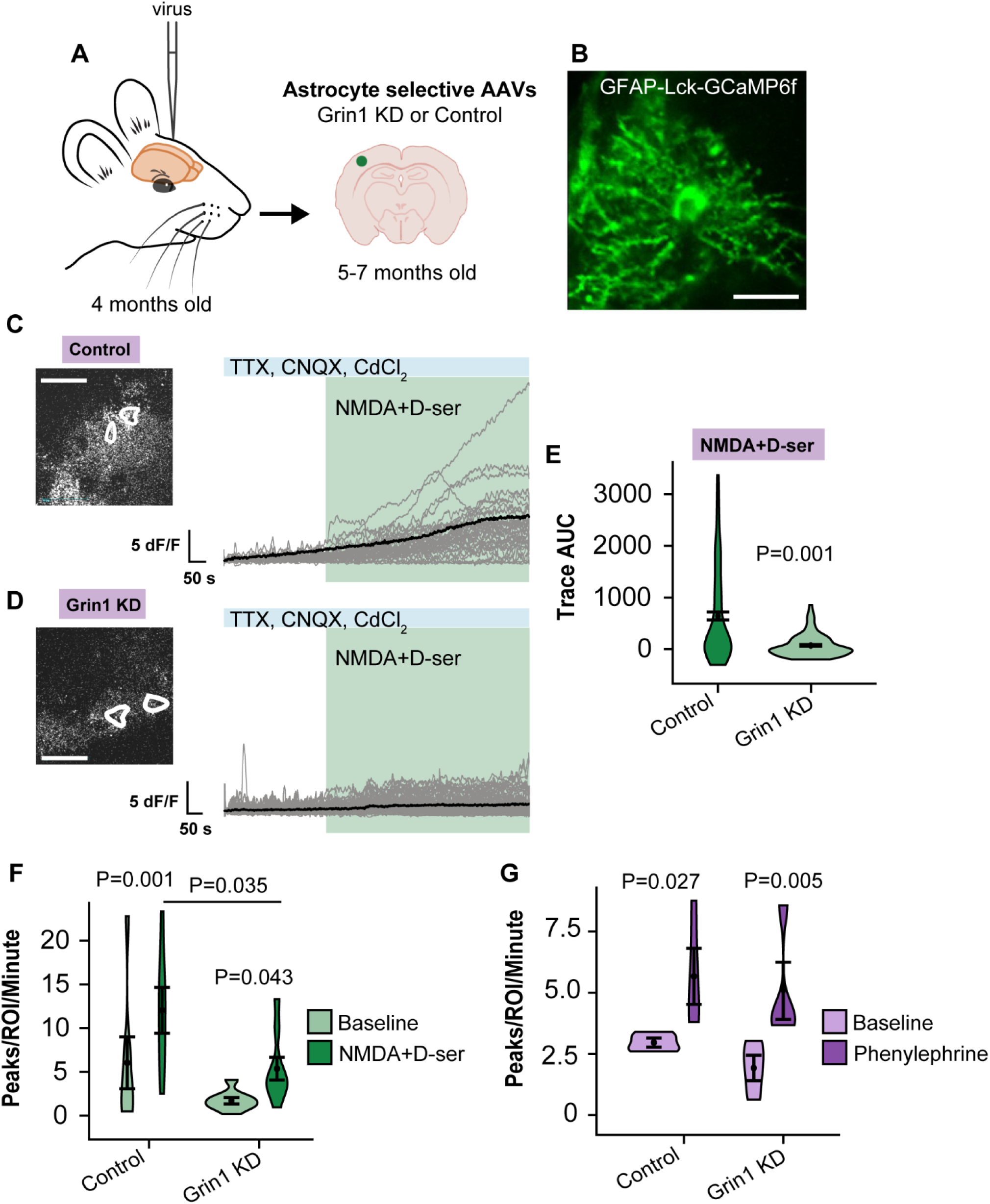
A functional reduction of astrocyte Ca^2^ responses to NMDAR agonists after Grinl KD. A) Experimental schematic for GrinlKD or control virus injections and brain slices. B) Representative brain slice astrocyte image. (Scale= 20 µm) C, D) Left: White shapes are example astrocyte regions of interest (ROIs) with Ca^2+^ events during agonist application (NMDA (50 µM) + D-serine (10 µM)) from Grin1 KD or control brain slices. (Scale= 50 µm). Right: individual ROI Ca^2+^ traces (grey) with the mean Ca^2+^ response (black) to agonist application in the presence of neuronal blockers. E) Area under the curve (AUC) of traces evoked by NMDA + D-serine. F) Number of Ca^2+^ peaks per ROI per minute evoked by NMDA + D-serine. Control: n= 165 ROIs from 8 mice, Grin1 KD: n= 136 ROIs from 7 mice. G) Number of Ca^2+^ peaks per ROI per minute evoked by phenylephrine (10 µM). Control: n= 50 ROIs from 5 mice, Grin1 KD: n= 51 ROIs from 5 mice. All statistics were calculated using linear mixed models and Tukey post hoc tests.

### Astrocyte NMDAR knockdown reduces sensory-evoked microdomain calcium events *in vivo*

To investigate the impact of Grin1 KD on Ca^2+^ events in cortical astrocytes and neurons *in vivo*, we prepared a chronic cranial window over the whisker barrel cortex in adult mice injected with a combination of virus for astrocyte control or Grin1 knockdown with AAV for neuronal expression of red GECI RCaMP1.07 (AAV9-hSYN-RCaMP1.07; Fig. 3A). Following recovery from surgery and mapping of the barrel cortex, mice were trained for awake two-photon imaging ^10^ to capture cortical Layer 2/3 (L2/3) astrocyte and neuron Ca^2+^ signaling during individual whisker vibrations (90Hz, 8s) corresponding to the virus injection area (Fig. 3A, B, and D). Calcium event ROIs were manually selected (for neuronal somata) or identified based on their activity (astrocyte microdomains and neuronal dendritric events) using the MATLAB toolbox, CHIPS (Fig. 3C; ^10,46,47^). Due to the membrane localization of the sensor, astrocyte Lck-GCaMP6f Ca^2+^ events occurred as spatially-confined ROIs throughout the “astropil” (i.e. Ca^2+^ MDs). Both control and Grin1 KD astrocytes had a similar number of spontaneous MDs per field of view (ROIs/FOV) in trials without whisker stimulation (mean ± SEM; 2.01 ± 0.12 control vs. 2.26 ± 0.17 KD ROIs/FOV; Fig. 3E). As shown previously ^10,13,15^, whisker stimulation evoked more MDs in control astrocytes (3.21 ± 0.23 ROIs/FOV) and higher Ca^2+^ event amplitudes (1.93 ± 0.05 dF/F; Fig. 3E, F). However, in Grin1 KD astrocytes, whisker stimulation failed to increase the number of MDs (1.93 ± 0.12 ROIs/FOV) or Ca^2+^ amplitudes (1.99 ± 0.06 dF/F; Fig. 3E, F; Supplementary Fig. 2). The nature of the sensory stimulus did not influence this reduction in Grin1 KD astrocyte MDs, since strong electrical stimulation (400 µA, 4Hz for 5 s) applied to the whisker pad of anesthetized mice also failed to increase the number of MDs in Grin1 KD astrocytes (Supplementary Fig. 3A, B). During whisker stimulation, astrocyte MDs in Grin1 KD astrocytes covered a smaller area, since the fraction of active astrocyte pixels relative to the total astrocyte area was lower (0.07 ± 0.007 control vs. 0.01 ± 0.002 KD; Fig. 3G, H). Furthermore, the fraction of astrocyte area that was active in more than one stimulation trial was lower in Grin1 KD astrocytes (repeated response score; 0.060 ± 0.005 control vs. 0.036 ± 0.008 KD; Fig. 3G, I), suggesting that fewer MDs were repetitively activated across multiple stimulation trials compared to control astrocytes.

**Figure 3.**
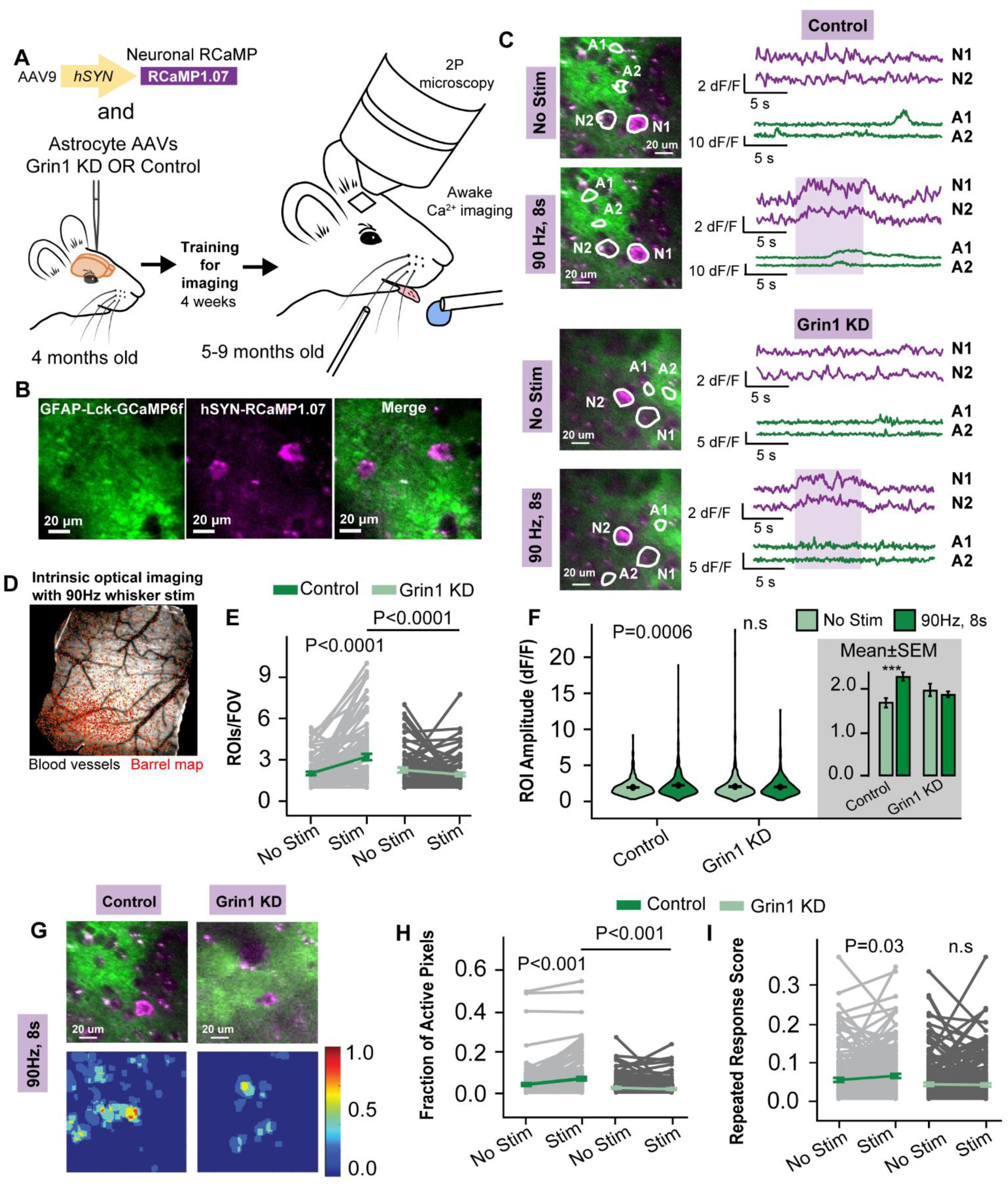
A reduction of stimulation-evoked Ca^2^ events in Grin1 KD astrocytes *in vivo.* A) Experimental schematic for awake Ca^2+^ imaging of neurons (RCaMP1.07) and Grin1KD or control astrocytes (Lck-GCaMP6f) after AAV injection and cranial window surgery. B) Representative astrocytic Lck-GCaMP6f and neuronal RCaMP1.07 expression. C) Example neuron (purple) and astrocyte (green) ROIs (left) and their traces (right) during trials with no stimulation or whisker stimulation (90 Hz, 8 s, purple shaded area). D) Example whisker barrel map (red) through the cranial window during intrinsic optical imaging. Virus expression was confirmed to be localized within this area for 2P imaging. E) The number of astrocyte microdomain ROIs per field of view (FOV) was reduced in Grin1 KD during stimulation. F) The mean astrocyte ROI Ca^2+^ amplitude was reduced in Grin1 KD during stimulation. G) Example response heatmaps (bottom) for control and Grin1 KD astrocytes (top) generated by overlaying microdomain ROI masks from all trials with stimulation. The scale was the fraction of trials with a response in that area. H) The fraction of active astrocyte pixels (due to ROI activity) per FOV was reduced during stimulation in Grin1 KD mice. I) The fraction of astrocyte pixels active in two or more trials (repeated response score) increased with whisker stimulation in control and not Grin1 KD mice. Control: n= 1127 ROIs from 103 FOV in 8 mice, Grin1 KD: n= 448 ROIs from 97 FOV from 11 mice. Comparisons made by linear mixed models and Tukey post hoc tests. *P < 0.05, **P < 0.01, and ***P < 0.001.

### Astrocyte NMDAR knockdown impairs neuronal activity *in vivo*

We also measured RCaMP1.07 Ca^2+^ events as a proxy for neuronal activity in the cortical circuit surrounded by astrocytes with reduced NMDAR expression. Spontaneous neuronal Ca^2+^ events in trials without whisker stimulation occurred more frequently (2.62 ± 0.24 control vs. 3.84 ± 0.32 KD neurons and dendrites per field of view i.e. ROIs/FOV) and with higher amplitude (1.23 ± 0.03 control vs. 1.70 ± 0.09 KD vs. dF/F) near Grin1 KD astrocytes compared to controls (Fig. 4A, B), suggesting that the spontaneous neuronal activity was elevated. Upon whisker stimulation, more active neurons and dendrites were detected in control mice (4.77 ± 0.43 ROIs/FOV), but the number of active neurons near Grin1 KD astrocytes did not change (3.13 ± 0.16 ROIs/FOV; Fig. 4C), indicating a reduced recruitment of responding neurons during cortical circuit activation. A similar reduced neuronal recruitment was observed following an electrical whisker pad stimulation (Supplementary Fig. 3C), suggesting that strong stimuli do not overcome neuronal impairments following astrocyte Grin1 KD. Furthermore, these effects were not influenced by sex, as the responses and effect of astrocyte Grin1 KD were the same for neurons and astrocytes from males and females (Supplementary Fig. 4). Although there was an overall decrease in the number of responding neuron ROIs near Grin1 KD astrocytes, the mean amplitude of the stimulation-evoked Ca^2+^ events in the remaining responding neurons was larger than controls (1.25 ± 0.02 KD vs. 1.09 ± 0.01 control dF/F; Fig. 4D).

**Figure 4.**
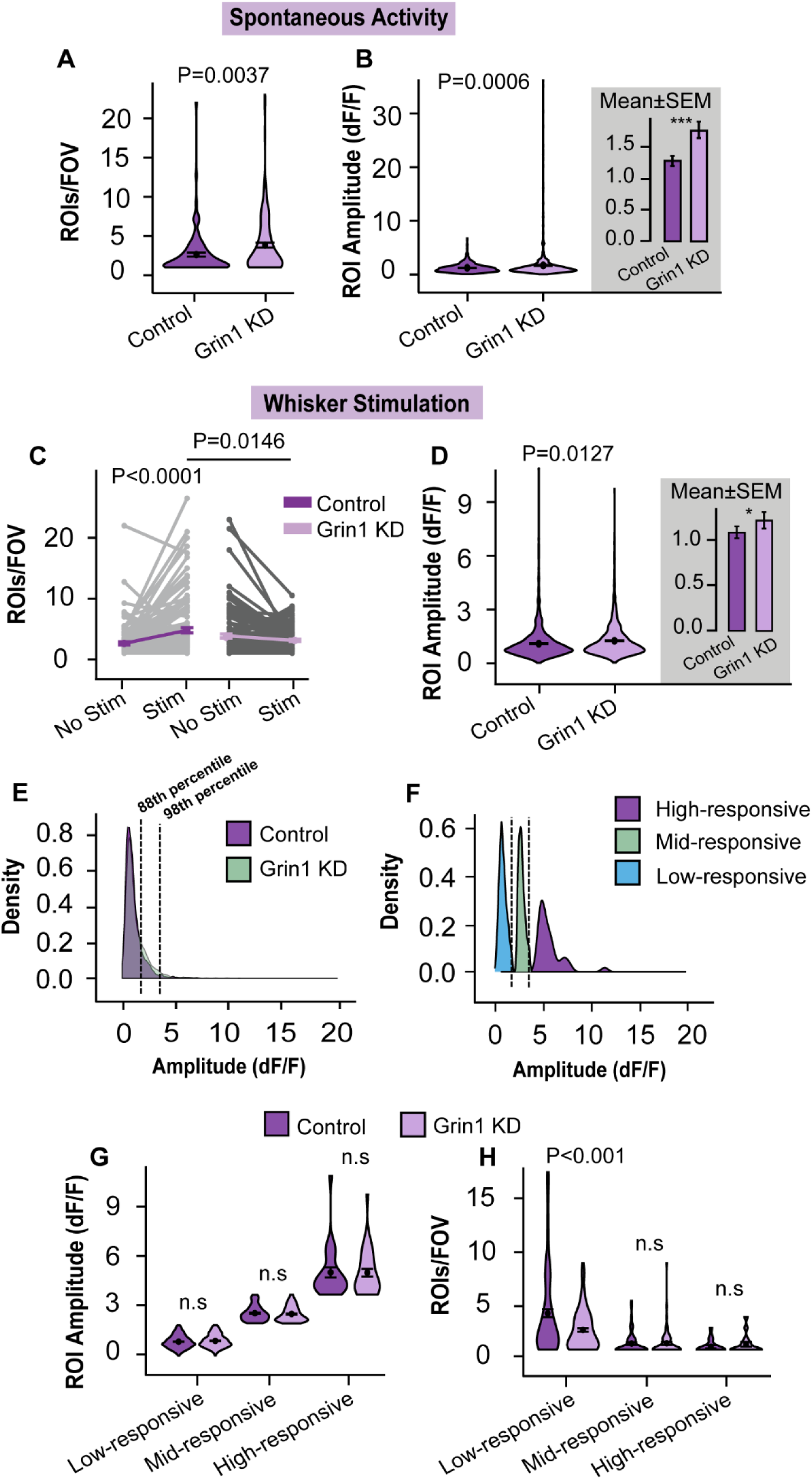
Altered neuronal activity after astrocyte Grin1 KD *in vivo.* A) The number of spontaneous neurons and dendrites per field of view increased in Grin1 KD mice (ROIs/FOV). B) ROI Ca^2+^ amplitude in spontaneous neurons was increased in Grin1 KD mice. C) Whisker stimulation (90Hz, 8s) failed to increase the number of active neurons (ROIs) per FOV in Grin1 KD. D) ROI Ca^2+^ amplitude was elevated during stimulation in Grin1 KD. Control: n=1799 ROIs from 122 FOV in 8 mice, Grin1 KD: 1210 ROIs from 144 FOV in 11 mice. E) Distribution of neuronal ROI amplitudes (dF/F) for control and Grin1 KD. Left dashed line is the 88^th^ percentile (1.92 dF/F) and the right dashed line is the 98^th^ percentile (3.47 dF/F). F) Classification of neuron types based on amplitude percentiles in E: high responding neurons, mid responding neurons, and low responding neurons. G) ROI Ca^2+^ amplitudes for each class of neuron. H) Number of each neuronal type per FOV showed that there are fewer low amplitude neurons in Grin1 KD. Control, High-responsive: n= 35 ROIs from 20 FOV, Mid-responsive: n= 158 ROIs from 58 FOV, Low-responsive: n= 1606 ROIs from 121 FOV, 8 mice. Grin1 KD, High-responsive: n= 37 ROIs from 19 FOV, Mid-responsive: n= 164 ROIs from 69 FOV, Low-responsive: n= 1009 ROIs from 141 FOV, 11 mice. Comparisons made by linear mixed models and Tukey post hoc tests. ***P < 0.001.

Previous studies have classified neurons based on the amplitude of their Ca^2+^ events into low, mid and high responsive cells ^48^, which reflects their level of recruitment during sensory stimulation. High-responsive neurons have reliable, large amplitude Ca^2+^ events during whisker vibrations ^48,49^, and this small population of cells “sparsely” encodes sensory stimulation ^49–53^. Low and mid-responsive neurons are more weakly recruited during repeated trials of sensory stimulation, but they have a greater capacity for reorganization during changes in sensory input ^48^. In order to consider sparse encoding and different neuronal populations, we classified neurons in both Grin1 and control mice based on the average neuronal Ca^2+^ amplitude during stimulation, as low-responsive (amplitude< 88^th^ percentile of 1.92 dF/F), mid-responsive (amplitude between 99^th^ percentile of 1.92 dF/F and 98^th^ percentile of 3.47 dF/F) or high-responsive (amplitude > 98^th^ percentile of 3.47 dF/F; Fig. 4E, F). Overall, the calcium amplitudes were the same for each class of neurons near both control and Grin1 KD astrocytes (Fig. 4G), even though the pooled average amplitude for all ROIs was greater near Grin1 KD astrocytes (Fig. 4D). Interestingly, the number of low-responsive neurons was reduced in Grin1 KD mice (4.37 ± 0.37 control vs 2.80 ± 0.15 KD ROIs/FOV, Fig. 4H), which would account for the decreased number of ROIs/FOV in the total population (Fig. 4C). Sparse coding of the whisker stimulus was still intact because the number of high-responsive cells was the same between controls and Grin1 KD (1.34 ± 0.16 vs 1.41 ± 0.18 ROIs/FOV; Fig. 4G), which could explain the elevated average amplitude from the total neuron population in Grin1 KD mice (Fig. 4D).

Neurons in the whisker barrel cortex increase their levels of synchronicity during stimulation ^49,54,55^. In order to estimate synchronization, we calculated the Pearson’s correlation coefficient between pairs of neurons in the same field of view in each trial. Improved synchronization occurred during stimulation in neurons near control astrocytes, as the correlation increased during trials with whisker stimulation compared to trials with no stimulus (−0.01 ± 0.002 no stim vs 0.03 ± 0.001 stim; Fig. 5A). For Grin1 KD, the neuronal correlation decreased during stimulation (0.06 ± 0.001 no stim vs 0.008 ± −0.01 stim; Fig. 5A). These effects were most pronounced between pairs of high-responsive neurons, as their synchronicity increased greatly during whisker stimulation near control astrocytes (−0.39 ± 0.003 no stim vs 0.37 ± 0.02 stim; Fig. 5B). However, near Grin1 KD astrocytes, high-responsive neurons had a greater correlation during trials without stimulation (0.68 ± 0.003), reflecting higher spontaneous activity, but the correlation dropped during stimulation (−0.16 ± 0.02; Fig. 5B). This suggests that reduced synchronicity that may impact sparse coding and network function following astrocyte Grin1 KD.

**Figure 5.**
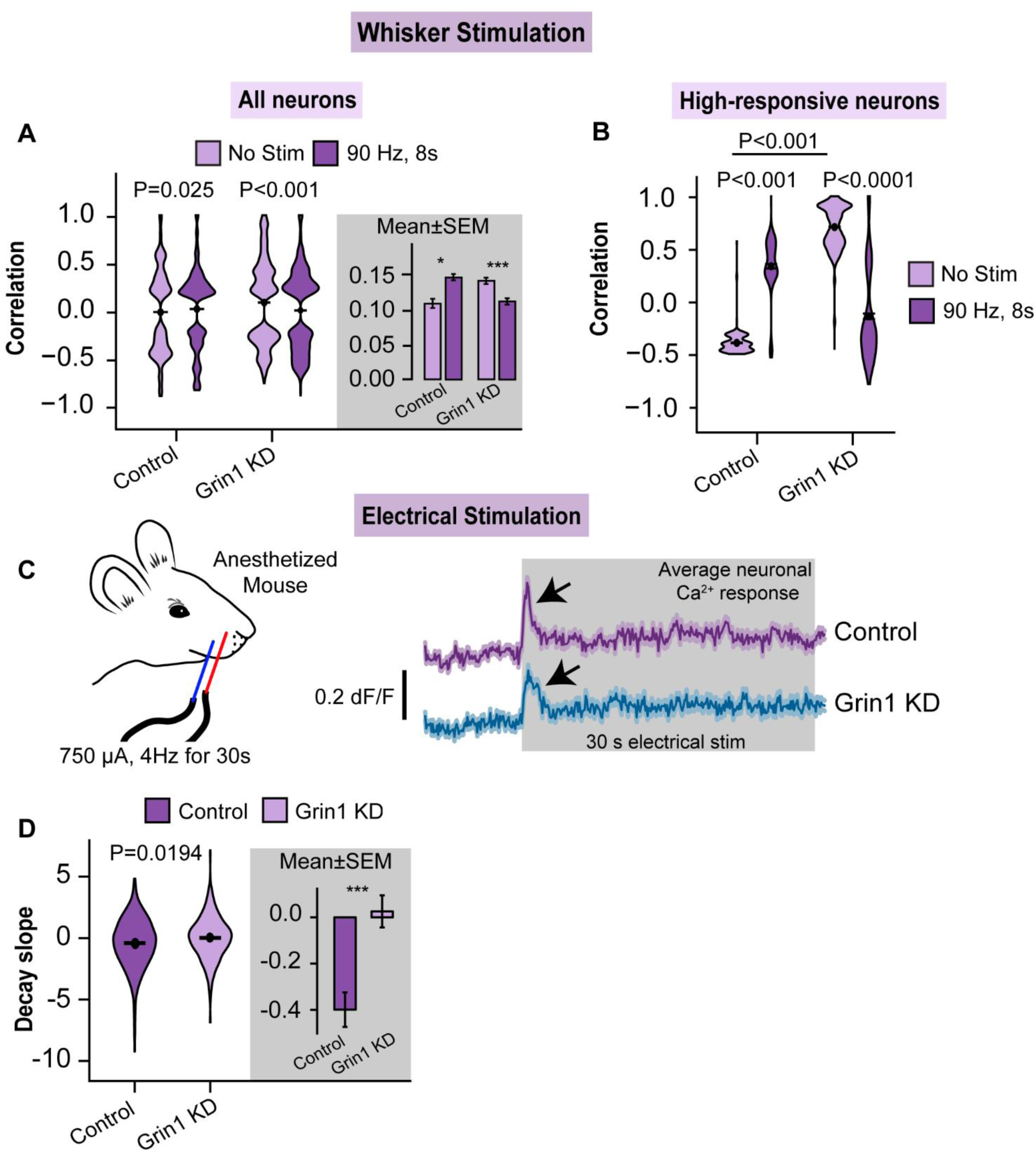
Alterations in neuronal synchronization and adaptation. A) Neuronal synchronization was determined by Pearson’s correlation between stimulation-activated neurons (i.e. Ca^2+^ event during the stimulus period (8s)) from the same FOV. Overall neuronal synchrony increased with stimulation in control and not in Grin1 KD. Control: n= 94167 comparisons from 8 mice; Grin1 KD: n=40376 comparisons from 11 mice. B) Neuronal correlation within the high-responsive population showed that high-responsive neurons were less synchronized in Grin1 KD during whisker stimulation (Control, No Stim: n= 1175 comparisons from 15 ROIs, Stim 90 Hz: n= 222 comparisons from 31 ROIs, 8 mice). (Grin1 KD, No Stim: n= 3266 comparisons from 62 ROIs, Stim 90 Hz: n= 367 comparisons from 36 ROIs, 11 mice). C) Prolonged electrical stimulation of the whisker pad (750 µA, 4Hz for 30s) resulted in robust neuronal Ca^2+^ events that quickly adapt (black arrows) for the duration of the stimulus. D) The slope of the Ca^2+^ event decay was calculated from time 0.5-1 s after the start of the stimulus. The adaptation was slower in Grin1 KD mice. Control: n=714 ROIs from 9 mice; Grin1 KD: n=629 ROIs from 8 mice. Data represented as mean ± SEM. All statistics were calculated using linear mixed model and Tukey post hoc tests.

Neurons adapt their response during prolonged periods of activity, such as long sensory stimulation. We applied an electrical stimulus to the whisker pad (750 µA, 4Hz) for prolonged periods (30s) in anesthetized mice transduced with control or Grin1 KD viruses and neuronal RCaMP1.07. Neurons in both control and Grin1 KD mice responded to the stimulus with a sharp Ca^2+^ peak that adapted within a few seconds of the start of stimulation (Fig. 5C). We calculated the slope of this signal decay in Ca^2+^ amplitude and found that control neurons had a more negative decay slope compared to Grin1 KD neurons (Fig. 5D), suggesting that they adapted more quickly. This is the first evidence that changes to astrocytic Ca^2+^ MD signaling can impact and delay neuronal adaptation.

### Astrocyte NMDAR knockdown reduces fast-onset Ca^2+^ microdomains

The majority of astrocyte Ca^2+^ MDs evoked by sensory stimulation have a delayed signal onset (~5s after the start of the stimulus ^10,56,57^) and many of these events are evoked by IP_3_R2 signaling and neuromodulators ^10^. A population of fast-onset astrocyte Ca^2+^ MDs that highly correlate with the onset of neuronal activity has also been identified ^10,15,58^, but the mechanism that underlies these fast signals has not been established ^10^. To determine if astrocyte NMDAR activity influences the onset of Ca^2+^ events, we considered the onset latency of neuron and astrocyte Ca^2+^ signals in control and Grin1 KD mice (Fig. 6A). First, the mean onset latency for neurons that responded during whisker stimulation near both control and Grin1 KD astrocytes was similar (2.53 ± 0.03 control vs 2.60 ± 0.04 s KD; Fig. 6B). Astrocyte Ca^2+^ MDs that occurred during whisker stimulation (0-8 s) were categorized as “fast” if their onset latency was less than the median onset latency for the neuronal population and “delayed” if their onset latency was slower than this value^10^. The number of fast MDs was reduced in Grin1 KD astrocytes compared to controls (Fig. 6C), demonstrating for the first time that ionotropic glutamate receptor signaling via astrocytic NMDAR can contribute to fast-onset Ca^2+^ events in astrocytes that potentially rapidly trigger the modulation of nearby synaptic activity. The number of delayed MDs evoked by stimulation was also lower in Grin1 KD astrocytes (Fig. 6C), suggesting that ionotropic glutamate receptors can also be activated on slower time-scales.

**Figure 6.**
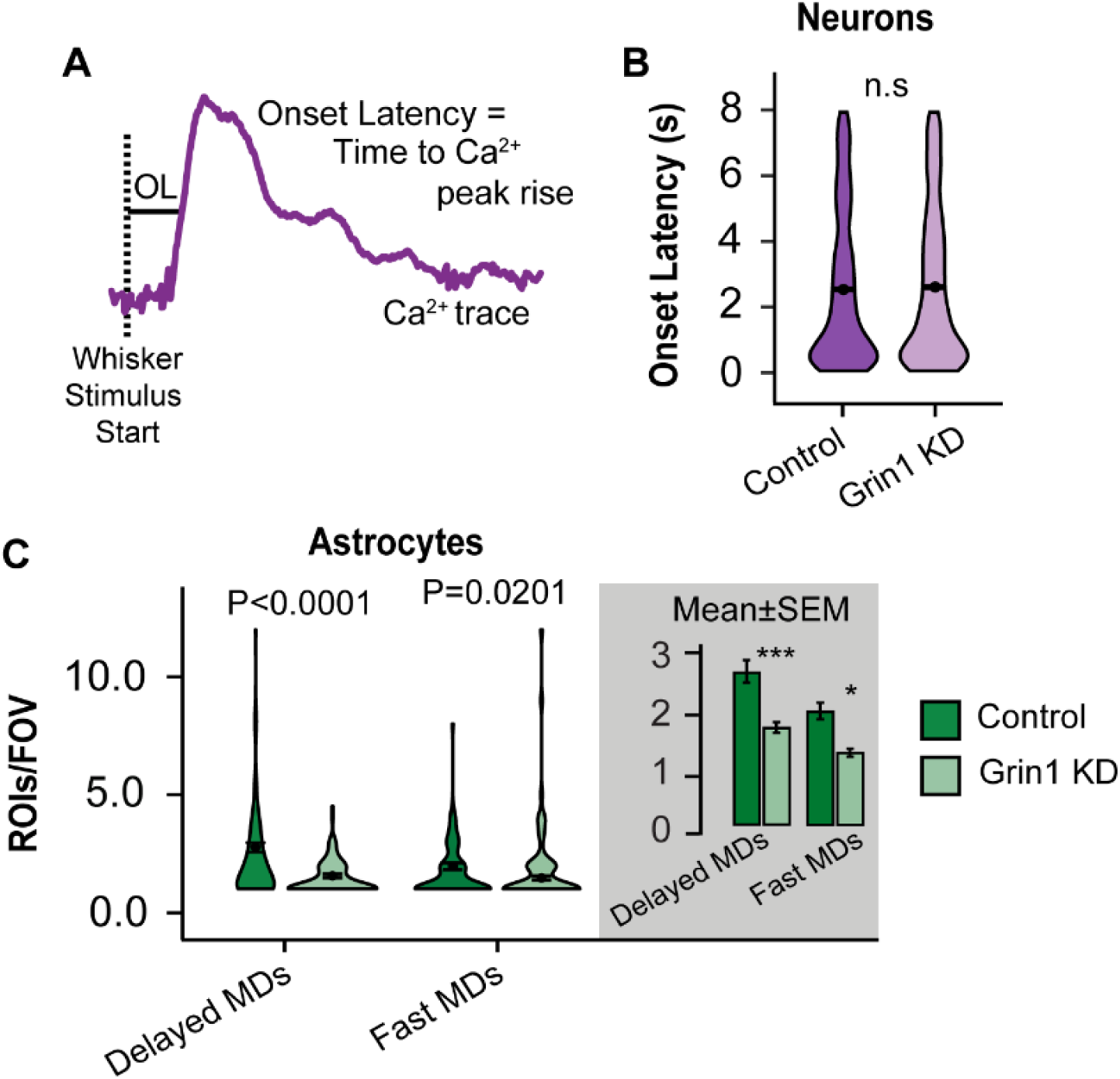
Rapid onset astrocyte Ca^2+^ microdomains are reduced in Grin1 KD. A) Onset latency (OL) is the earliest time point after the start of whisker stimulation at which the Ca^2+^ signal [dF/F] reached 2.5 SD of the baseline. B) The onset latency was not different between neurons in control and Grin1 KD neurons. Control: n=1799 ROIs, 8 mice, Grin1 KD: n= 1210 ROIs, 11 mice. C) Astrocyte Ca^2+^ microdomains (MDs) were classified as fast or delayed onset relative to the neuronal onset times. The number of fast onset and delayed onset MDs were reduced in Grin1 KD. Control, Delayed MDs: n= 796 ROIs from 103 FOV & Fast MDs: n= 331 ROIs from 76 FOV, from 8 mice; Grin1 KD, Delayed MDs: n= 240 ROIs from 78 FOV & Fast MDs: n= 193 ROIs from 56 FOV, from 11 mice. Data are represented as mean ± SEM. All statistics were calculated using linear mixed model and Tukey post hoc tests.

### Astrocyte NMDAR knockdown leads to sensory acuity impairments in behaviour

Given the impairment of neuronal recruitment we observed following astrocyte Grin1 KD, we tested the sensory perception of the animal using a whisker touch-based novel texture recognition test ^4,59^. In this task, following two days of acclimatization to the testing arena, the whiskers of the mice were trimmed such that only the whiskers corresponding to the virus injection area were long. Mice were habituated to two vertical objects fitted with identical sandpaper (e.g. 150 grit) and allowed to explore with their whiskers. One object was then replaced with an object covered in different sandpaper (e.g. 220 finer grit) of the same colour, and the time the animal spent exploring the textured objects was recorded over 3 min. Mice that can perceive the difference in textures will spend more time with the novel textured object, reflected as a positive discrimination index (difference in novel - familiar texture exploration time/ total exploration time). Mice with astrocyte Grin1 KD could not discriminate a small grit difference (ΔG = 32 µm; discrimination index near zero), but performed the same as controls when detecting larger grit differences (ΔG = 201 µm; Fig. 7B). This behaviour deficit was not due to a memory impairment because Grin1 KD and control mice performed similarly in the novel object recognition test (performed in the same manner as the novel texture recognition), where the objects are more distinct (different colours and shapes) and other cues are used to discriminate and remember objects (Fig. 7A, B).

**Figure 7.**
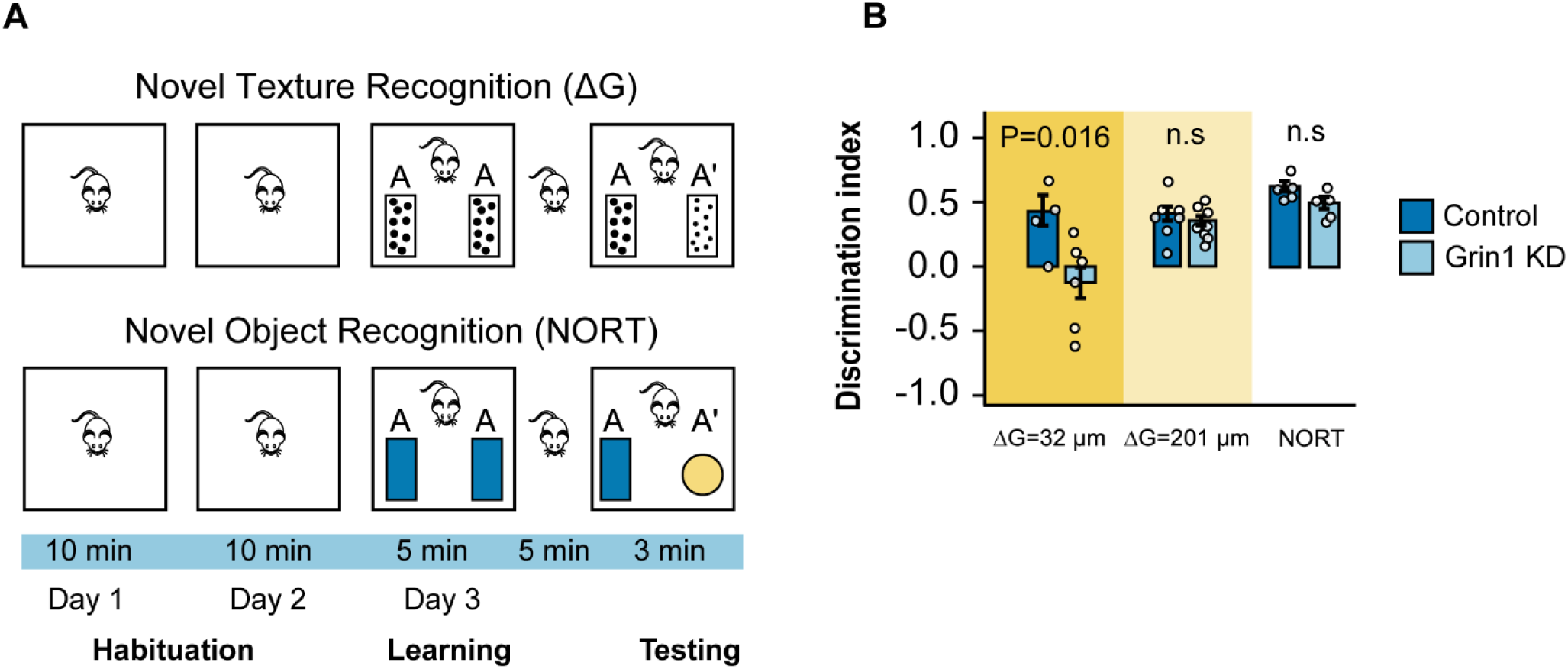
Sensory discrimination of the animal is reduced after astrocyte Grin1 KD. A) Schematic of whisker-mediated novel texture recognition (top) and novel object recognition test (bottom) tests. Mice ability to discriminate between different grades of sandpaper or two distinct objects is reported as discrimination index (difference in novel - familiar texture exploration time/ total exploration time) B) The discrimination index for small grit difference (ΔG= 32µm; 150 grit vs. 220 grit), large grit difference (ΔG= 200 µm; 60 grit vs. 220 grit) and the NORT. Circles=Individual animals. Comparisons were made with the Mann-Whitney-Wilcoxon test.

## Discussion

By utilizing a novel AAV-delivered shRNA knockdown approach to selectively reduce astrocyte NMDARs in the whisker barrel cortex, we show for the first time that astrocyte NMDARs contribute to rapid, stimulus-evoked microdomain Ca^2+^ events in awake mice. Astrocyte NMDARs are also linked to nearby cortical circuit function, since Grin1 KD reduced and desynchronized responses of neurons to the whisker stimulus and decreased neuronal adaptation. In fact, the effects of astrocyte NMDAR knockdown on the cortical circuit were significant enough to cause a sensory deficit in the animal. These findings represent a new direction for astrocyte-neuron interactions and a paradigm-shift for the mechanisms underlying astrocyte Ca^2+^ MDs away from second-messenger signaling and toward ionotropic mechanisms.

### Astrocyte Ca^2+^ microdomains and NMDAR signaling

Our work provides new knowledge about astrocyte NMDARs and their contribution to microdomain Ca^2+^ events. Previous evidence for the functionality of astrocyte NMDARs has been fraught with controversy because many early studies used primary astrocyte cultures from pups, which may have variable expression levels of NMDAR subunits (for review see ^29,32,60^). Also, many other brain cells express NMDARs (neurons, microglia, oligodendrocytes, endothelial cells, etc. ^61^), clouding the interpretation of results with bath-applied pharmacology on brain slices or mixed cultures. More recent studies have recorded astrocyte NMDAR-mediated currents or shown Ca^2+^ responses to NMDAR agonists in brains slices from adult mice ^26,31,36–38,40^. This has provided convincing support for the functional presence of these receptors, but it remained unclear if NMDAR activation induced localized Ca^2+^ MDs *in vivo*. We used a membrane-tagged genetically encoded calcium indicator (GCaMP6f) to detect Ca^2+^ fluctuations near the plasma membrane in adult mice at an age when NMDAR currents are known to be strongest (compared to juveniles; ^40^). Knockdown of astrocyte NMDARs did not affect the number of spontaneous Ca^2+^ events in astrocytes, but prevented the recruitment of Ca^2+^ MDs during whisker stimulation (Fig. 3). Synaptic activation can induce astrocyte NMDAR currents ^36^, but we provide the first evidence linking neuronal circuit activity, astrocyte NMDARs and Ca^2+^ microdomain events *in vivo.* This suggests that cortical astrocytes respond to synaptic glutamate via NMDAR activation and that this is an important component of how astrocytes encode sensory information.

Astrocytes express a plethora of GPCRs that can induce elevations in intracellular Ca^2+^ via IP_3_-mediated release of Ca^2+^ from intracellular stores. This signaling has been the focus of the astrocyte field, and now tools to exploit these pathways are used to activate (such as Gq-DREADDs) or inhibit (iβARK or IP_3_R2 knockout mice) astrocyte Ca^2+^ to help determine the functional roles of these cells ^2,5,62,63^. However, Gq-GPCR signaling is clearly not the sole contributor to astrocyte microdomain events, since IP_3_R2 knockout animals or mice expressing iβARK to disrupt the Gq pathway have remaining Ca^2+^ MDs in their processes ^5,10,12^. Furthermore, circuit activation via sensory stimulation can evoke fast-onset Ca^2+^ MDs in astrocytes that are independent of IP_3_R2 and Ca^2+^ release from internal stores ^10^. We theorized that these fast Ca^2+^ events could be mediated by extracellular calcium influx through ionotropic receptors ^60^, which is what led us to knockdown astrocyte NMDAR. Indeed, we found a reduction in the number of fast-onset MD Ca^2+^ signals in astrocytes that were on a similar timescale to neurons (Fig. 6), confirming that ionotropic receptors can contribute to rapid activity-based dynamics in astrocytes. This also suggests that these receptors could be important for triggering quick feedback mechanisms that alter synaptic activity. It is important to note that Ca^2+^ MDs were not completely abolished in Grin1 KD astrocytes (Fig. 3), and the number of Ca^2+^ events evoked by phenylephrine application to Grin1 KD slices was similar to controls (Fig. 2 G). This suggests that astrocyte Ca^2+^ and GPCR signaling were still intact.

It should be noted that multiple mechanisms may underlie NMDAR-mediated astrocyte Ca^2+^ MDs, through direct and indirect pathways. First, NMDARs are ion channels, so receptor activation could cause direct extracellular Ca^2+^ influx that would be readily visualized by our membrane-tagged GECI. Second, the influx of Na^+^ through NMDARs may elevate intracellular Na^+^ inducing reversal of the sodium-calcium exchanger and bringing in extracellular Ca^2+^ in exchange for Na^+^ ^26,35,64^. Third, ion influx through NMDARs may depolarize the astrocyte membrane, activating voltage-gated Ca^2+^ channels that permit extracellular Ca^2+^ influx ^27^. Finally, studies in cultured astrocytes have suggested the Ca^2+^ responses following NMDAR activation are at least partially mediated by Ca^2+^ release from intracellular stores ^65–68^. Conversely, astrocyte NMDARs may also promote the filling of intracellular Ca^2+^ stores during periods of strong, prolonged synaptic activity ^69^. Therefore, astrocyte NMDARs are potentially critical for regulating intracellular Ca^2+^ at multiple points from Ca^2+^ influx to store release. Since we observed a reduction in stimulation-evoked astrocyte MDs (Fig. 3) of both fast and delayed onset (Fig. 6) and delayed-onset Ca^2+^ events are known to be mediated by IP_3_R2 ^10^, it is tempting to speculate that both extracellular Ca^2+^ influx and interactions with internal Ca^2+^ stores are factors in astrocyte NMDAR signaling in the mouse cortical circuit. However, the precise Ca^2+^ mechanism(s) disrupted in astrocytes during our Grin1 KD approach remain to be determined.

### Cortical circuit regulation and sparse sensory coding

In L2/3 of the barrel cortex, 80% of neurons are excitatory pyramidal cells^51^ that are densely innervated by GABAergic interneurons ^51,70–72^. We used the hSYN promoter to drive RCaMP1.07 expression in all neurons (Fig. 4), but given that pyramidal neurons are the majority of the population, it stands to reason that we primarily measured excitatory cells. In quiet, awake animals, there is low pyramidal neuron activity and high activity of interneurons ^51,72–75^, which generates a blanket of inhibition that must be overcome during sensory processing ^76^. Whisker deflection evokes high contrast activity compared to baseline firing during quiet wakefulness ^72,77^, and opens up local transient “holes” in the blanket of inhibition. This is the basis for sparse encoding, where a group of active pyramidal neurons display large action potentials that can be detected as high amplitude Ca^2+^ events (high-responsive cells) ^76^. Thus, the balance of excitation and inhibition during quiet wakefulness and sensory stimulation improves the signal-to-noise ratio for reliable, efficient stimulus representation ^78^ and optimal sensory perception ^72^. We found that spontaneous neuronal activity during quiet wakefulness was higher near Grin1 KD astrocytes (Fig. 4A, B), and during whisker stimulation, fewer neurons were activated (Fig. 4C). These changes in neuronal activity were also reflected in the synchrony between neurons from the same field of view in Grin1 KD mice, where Ca^2+^ events were more synchronized during no stimulation trials, but less synchronized during whisker stimulation when compared to neurons in control mice (Fig. 5A, B). This suggests that astrocyte Grin1 KD disrupts the blanket of inhibition during quiet wakefulness increasing the “noise” at baseline. This would suppress the contrast or signal of whisker deflection-evoked responses needed for appropriate perception. Elevated spontaneous circuit activity or “noise” has been shown to impact animal performance in a discrimination test ^79^, and this could explain the reduced sensory acuity we observed in our novel texture recognition task following astrocyte Grin 1 KD (Fig. 7).

High-responsive neurons are the primary cells that encode sensory information ^49–53^. We found that the number of high-responsive cells in Grin1 KD mice was the same as control (Fig. 4H), suggesting that encoding of the whisker stimulus remained intact. However, high-responsive cells in Grin1 KD were less synchronized during whisker stimulation (Fig. 5B), which indicates network dysfunction and possible impacts on sensory processing. Beyond high-responsive cells, other nearby neurons also respond to whisker stimulation, but with low amplitude Ca^2+^ events and not in every stimulus trial. These low-responsive cells are recruited during plasticity ^48,50,80^ and play a role in recoding stimuli after a change in sensory experience ^48^. We found that the overall reduction in neuronal activity in response to whisker stimulation in Grin1 KD mice (Fig. 4C, D) was primarily mediated by a decreased number of low-responsive neurons since the number of high-responsive cells stayed the same (Fig. 4H). Although the contribution of low-responsive cells to network activity during sparse firing has been suggested to be marginal ^50^, our results indicate that they could be important for normal sensory processing, as their reduction was linked to decreased texture discrimination (Fig. 7). Furthermore, reduced recruitment of low-responsive cells following astrocyte Grin1 KD could be indicative of decreased plasticity ^48^. Both astrocyte Ca^2+^ signaling and astrocyte NMDARs have been linked to plasticity, since IP_3_R2 knockout mice have diminished cortical homeostatic plasticity^81^ and hippocampal astrocyte NMDARs mediate heterosynaptic plasticity^27,82^. Therefore, a reduction in cortical astrocyte NMDARs and Ca^2+^ MDs following our Grin1 KD approach may have effects on cortical plasticity, but further experiments are required to determine the influence of these pathways on this type of neuronal regulation.

Cortical neurons strongly adapt to prolonged trains of stimuli due to short-term depression at thalamocortical synapses ^83,84^. Adaptation is beneficial for sensory processing because the coding efficiency is enhanced permitting better discrimination of similar stimuli ^85–87^. When we applied prolonged electrical stimuli to the whisker pad of anesthetized control mice, neurons that responded to the stimulus had an adaptive decay in their Ca^2+^ events within the first few seconds of the start of stimulation (Fig. 5C, D). Neurons near Grin1 KD astrocytes adapted slower (prolonged decay slope), suggesting their adaptation response was impaired. Reduced adaptation could also contribute to the discrimination deficit that we observed in our novel texture recognition task (Fig. 7), since Grin1 KD mice had difficulty with more textures that were similar and adaptation enhances with discrimination of similar stimuli ^85–87^. At this stage, because our Grin1 KD virus is expressed across multiple cortical layers, it is not possible to determine if impairments in adaptation occur as a result of disrupted depression at thalamocortical synapses in deeper cortical layers or local changes in the L2/3 microcircuit.

Future directions will elucidate the underlying mechanism of astrocyte-neuron communication that caused elevated spontaneous neuron activity, reduced/unsynchronized responses to whisker stimulation, and sensory behaviour impairment after astrocyte Grin1 KD. An increase in spontaneous neuron activity could occur due to reduced tonic inhibition on pyramidal cells, suggesting that astrocytes are important contributors to cortical excitatory/inhibitory balance. This can be viewed as regulation of neural network gain control for optimal synchrony and information processing ^9^. In cultured cortical astrocytes, NMDAR agonists evoked ATP gliotransmitter release and this may alter phasic and tonic cortical inhibition in brain slices ^88^. Therefore, a loss of gliotransmission (particularly ATP) after astrocyte Grin1 KD may induce the changes we observed in cortical activity.

In conclusion, we demonstrate for the first time that astrocytes can influence mouse whisker barrel microcircuits needed for sensory acuity. We also provide novel evidence that astrocyte NMDARs are an essential component of cortical circuits. We propose that astrocytes rapidly respond to glutamatergic transmission with Ca^2+^ microdomains evoked by NMDAR activity. This triggers astrocyte-neuron feedback that regulates neuronal activity needed for accurate sensory processing. This is a novel direction for astrocyte Ca^2+^ signaling, away from GPCR pathways.

## Materials and Methods

### Mouse Models

All experimental procedures outlined below were approved by the Animal Care Committee of the University of Manitoba in accordance with the Canadian Council on Animal Care. Male and female C57BL/6NCrl mice were used in this study (Table S1). Animals were housed under a standard 12-hr light/dark cycle with *ad libitum* access to food and water (unless water was removed overnight for awake two-photon imaging and training). AAV viruses were injected at 3-5 months of age and data was collected between 5 and 9 months of age.

**Table S1.** Mouse sex and age used in the study.

| **Experiment** | **Number/Sex** | **Age at the time of surgery** | **Age at the time of experiment** |
| --- | --- | --- | --- |
| FACS and qPCR | 10 Control, 9 Grin1 KD All Female | 4-5 months | 5-6 months |
| Immunohistochemistry | 6 Control, 6 Grin1 KD<br>3 Male and 3 Female | 4 months | 6 months |
| Brain slice Ca<sup>2+</sup> imaging | 5 Control, 7 Grin1 KD<br>All Female | 5 or 7 months old | 7 or 9 months old |
| *In vivo* awake Ca<sup>2+</sup> imaging | 8 Control; 4 Female and 4 Male<br>11 Grin1 KD; 6 Female and 5 Male | 3-5 months | 3-5 months |
| Whisker-mediated discrimination task | 4 Control, 5 Grin1 KD<br>All female | 3-5 months | 7 months |
| Novel object recognition task | 5 Control; 3 Female and 2 Male/<br>5 Grin1 KD; 2 Female and 3 Male | 4 months | 6 months |

### Strategy for Grin1 knockdown and virus production

The AAV9-hSYN-RCaMP1.07 was packaged by the Vector Core at the University of North Carolina at Chapel Hill and has been used previously *in vivo* ^10^. The shRNA^mir^ silencing and non-silencing constructs were custom designed. Multiple online siRNA prediction sites were used to search for optimized targeting sequences within the mouse Grin1 coding sequence to identify target regions that were consistent between different algorithms. In the end, 3 target regions, 5’-GTTGAGCTGTATCTTCCAAGAG-3’, 5’-CTATAGTTGGCAAACTTCCGGT-3’, 5’-CTTGATGAGCAGGTCAACGCAG-3’, were chosen that include the 5’ UTR, the ORF and the 3’ UTR. The 19nt targets were extended 5’ and 3’ to a final length of 22 nucleotides and modified to a sense and antisense miR-30-based shRNA sequence. A non-silencing hairpin derived from the pGIPZ library was used as our control. All sequences were cloned *in silico* into the optimized MIR-E backbone ^89^, and subsequently linked together to form a chain of 3 hairpins, separated by a spacer sequence. This shRNAmir-e multimer was encoded within a chimeric intron (chI) sequence between the splice donor and splice acceptor branch point sites. This entire chI-[3X(shRNAmir-e)] cassette, along with cloning sites, was synthesized using the ThermoFisher GeneArt Gene Synthesis service. The shRNAmir-e cassettes were digested out of the GeneArt constructs and cloned into a pssAAV-2-sGFAP-Lck-GCaMP6f backbone, generating the pssAAV-2-sGFAP-chI-[3X)shRNAmir-e)]-Lck-GCaMP6f. Ligated constructs were transformed into MDS42 E. coli cells by heat shock and grown in Terrific Broth (TB) + carbenicillin with vigorous shaking at 37°C for 16 hours. Plasmids were isolated using the PureLink HiPure Plasmid Maxiprep kit (ThermoFisher Scientific), eluted in sterile water, and sequence was confirmed (3’ end of sGFAP promoter, entire hairpin cassette, and Lck-GCaMP6f). These silencing and non-silencing constructs were packaged in AAV9; hGFAP-chI[3x(shmGrin1)]-Lck-GCaMP6f and hGFAP-chI[3x(shm/rNS)]-Lck-GCaMP6f, by Viral Vector Facility, University of Zurich.

### Headpost-implantation surgery

The cranial window surgery was conducted as previously described ^10^, in two separate surgeries 48-72 hours apart. First, animals were anesthetized with isoflurane (4% induction, %1.5-2 maintenance) and were fixed in a stereotaxic frame (RWD; Model 68507). A headcap was fitted on the skull using one layer of bonding agent (Bisco Dental; All Bond Universal Adhesive) and a few layers of dental cement (Ivoclar; Tectric EVOFlow) polymerized with blue light. A custom-made aluminum head post was attached to the headcap at the back of the head. The headcap covered all areas of the skull except for the left somatosensory cortex which was later used for craniotomy and virus injection. Animals were given 0.3 ml 5% glucose to help recovery. Animals also received Meloxicam (2mg/Kg; Metacam) every 24 hours for several days after the surgery until recovery, and Buprenorphine Slow Release (SR;0.5 mg/Kg) every 72 hours over 6 days.

### Intrinsic optical imaging

Two days after headpost implantation surgery, the somatosensory cortex was mapped using intrinsic optical imaging (IOI) through skull to localize specific whisker areas and identify proper regions for virus injection. Animals were anesthetized with isoflurane (4% induction and 0.5-1% maintenance) and head-fixed using the implanted headpost. The skull was washed with sterilized cortex buffer (NaCl, 125 mM; KCl, 5 mM; glucose, 10 mM; HEPES, 10 mM; CaCl_2_, 2 mM; MgSO4, 2 mM; pH~7.4) to visualize the vasculature. The solution was then replaced with ultrasound gel (HealthCare Plus) and a small cover slip (5 mm diameter; Fisher Scientific) was placed on top. Images were acquired using a 12-bit CMOS camera (Basler Ace acA2040-55µm) focused 400 µm below the cortical surface, under 630 nm red light illumination. Whisker stimulation (90Hz, 10 s), by lateral deflection of a single whisker threaded into a glass capillary affixed to a piezo actuator (PiezoDrive), increased the blood flow to the corresponding area of the somatosensory cortex which was identified by increased light absorption.

### Virus-injection surgery

On the second step of the surgery, following IOI, animals were anesthetized with triple-anesthesia including fentanyl (0.05mg/kg), medetomidine (0.5mg/kg), and midazolam (5mg/kg) injected subcutaneously. A craniotomy was cut over the barrel cortex. According to the IOI map, using a glass micropipette (Sutter Instruments; P97), and a custom-made hydraulic pump, 300 nL of virus mix was injected at 50 nL/min rate at a depth of 400 µm into the responding whisker areas. A mixture of AAV9-sGFAP-shRNA-Lck-GCaMP6f (1×10^12^ particles/ml) and AAV9-hSYN-RCaMP1.07 (2.4×10^12^ particles/ml) for Grin1 KD or AAV9-sGFAP-NS-shRNA-Lck-GCaMP6f (1×10^12^ particles/ml) and AAV9-hSYN-RCaMP1.07 (2.4×10^12^ particles/ml) for control animals. A square sapphire glass (3×3 mm) was lightly pressed on the exposed brain using a stereotaxic arm and fixed with dental cement to the head cap. Animals were given 0.3 ml 5% glucose to help recovery, and anesthesia antagonist (flumazenil and atipamezole, 0.5mg/kg and 2.5 mg/kg, respectively). Animals received Meloxicam (2mg/Kg) every 24 hours for several days after the surgery until complete recovery. Mapping of the barrel cortex (IOI) was repeated through the cranial window two weeks after surgery and before two-photon imaging commenced.

### Behavior training for awake imaging

One week after surgery, training started for awake two-photon imaging. Animals were handled twice a day for 3-5 days until they were comfortable being handled by the experimenter. Then, they were introduced to the head fixation multiple times a day for 3-4 days by restraint inside a custom apparatus tube with a holder for the implanted head post. Restraint started from several seconds and increased to several minutes as the animal became accustomed to the setup. Finally, when animals were acclimatized to the head fixation tube, they were water-deprived overnight and they were presented with water from a lick spout while restrained and their whiskers were periodically stimulated. Starting with short trials (12.5 seconds), a water drop was presented in simultaneously with an auditory cue at the end of each trial (10^th^ second). The aim was to train the animal to sit still, accept whisker stimulus during the trial, and receive water as a reward. The whisker stimulus was presented by threading a single whisker into a glass capillary affixed to a piezo element (PiezoDrive) vibrated at 90 Hz. The length and the number of trials were increased to 25 s and up to 50 trials per session, as the animal showed signs of being accustomed to the setup. Animals were trained two sessions a day, 3-5 days a week, for a total time of 2-4 weeks depending on their performance.

### *In vivo* two-photon Ca^2+^ imaging

Awake, water-deprived animals were imaged while head-restrained in the water-reward task setup under a two-photon laser-scanning microscope (Ultima In Vivo, Bruker Fluorescence Microscopy) with a 20x water immersion objective (N20X-PFH 20X Olympus XLUMPLFLN Objective, 1.00 NA, 2.0 mm WD). RCaMP1.07 and Lck-GCaMP6f were excited at 990 nm with a Ti:sapphire laser (Coherent, Ultra II) and emission light was split with a 565 LP dichroic to GaAsP photomultiplier tubes with 595/50 nm band pass filter (red) or a 525/70 band pass filter (green). At the start of each imaging session, fields of view in cortical layer 2/3 (depth 110-280 µm) were identified based on fluorescence expression and the corresponding IOI barrel map for each whisker. Short trials (12.5 s) with 3 s of whisker stimulation (90Hz) were used to confirm that neurons responded to whisker stimulation and that the correct whisker was stimulated. Once the cells were selected, a high resolution (512×512 pixels, 1.17 fps) image was collected of the field of view as a reference (to avoid imaging same fields of view across multiple sessions). Then, data was acquired at 128*128 pixels (13.84 fps) for 25-second trials including 5 seconds of baseline, 5 seconds of whisker stimulation (90 Hz) followed by 12 seconds without stimulation. For each field of view, a total of 10 trials were performed including 5 trials without stimulation alternating with 5 stimulation trials. The whisker stimulus was presented by threading whiskers into a glass capillary affixed to a piezo element vibrated at 90 Hz. Animals were imaged 3-5 days a week for up to 2 months.

For experiments with electrical whisker pad stimulation, animals were anesthetized with isoflurane (1.5%) and two electrodes were implanted superficially in the skin within the whisker area. Mild electric stimuli (4Hz, 1 ms pulse every 249 ms) were applied at 400 or 750 µA from 5 or 30 s. Images on the two-photon microscope were acquired in the same manner as awake imaging above.

### Brain slice two-photon Ca^2+^ imaging

Animals were anesthetized with isoflurane, cervically dislocated and decapitated. Brain was removed and rapidly placed in ice-cold carbogen-saturated (Oxygen, 95%; Carbon dioxide, 5%) slicing buffer (N-methyl-D-Glucamine, 93 mM; KCl, 3 mM; MgCl_2_ * 6H2O, 5 mM; CaCl_2_ * 2H_2_O, 0.5 mM; NaH_2_PO_4_, 1.25 mM; NaHCO_3_, 30 mM; HEPES, 20 mM; Glucose, 25 mM; Sodium Ascorbate, 5 mM; Sodium Pyruvate, 3 mM). The hemisphere injected with virus was cut and mounted on a vibratome with slicing chamber containing ice-cold oxygenated slicing buffer. Sagittal slices of 300 µm thickness were cut (50 Hz, 1.25 Amp) and incubated in oxygenated 32°C recovery solution (NaCl, 95 mM; KCl, 3 mM; MgCl_2_ * 6H2O, 1.3 mM; CaCl_2_ * 2H2O, 2.6 mM; NaH_2_PO_4_, 1.25 mM; NaHCO_3_, 30 mM; HEPES, 20 mM; Glucose, 25 mM; Sodium Ascorbate, 5 mM; Sodium Pyruvate, 3 mM) for 1 hour.

Astrocyte Ca^2+^ events were recorded in oxygenated aCSF (NaCl, 125 mM; KCL, 2.5 mM; NaH_2_PO_4_, 1.25 mM; MgCl_2_, 1 mM; CaCl_2_, 2 mM; NaHCO_3_, 25 mM; glucose, 25 mM) at 35°C using Ultima In Vitro Multiphoton Microscope (Bruker Fluorescence Microscopy) with a 40x water immersion objective (Olympus). Lck-GCaMP6f was excited at 930 nm and a high resolution (512*512 pixels) scanning of the slice was done to identify Lck-GCaMP6f-expressing astrocytes. Once the recording area was determined, a cocktail of neuronal activity blockers including TTX (1 µM), CNQX (10 µM), and CdCl_2_ (100 µM) was bath-applied to the slice for 20 minutes. Then, images (256×256 pixels, 2.58 frames per second (fps)) were recorded for 15 minutes while NMDA (50 µM) and D-serine (10 µM) were applied for the first 5 minutes and then washed out with aCSF for the next 10 minutes. After washing for 20 min, another 15-minute image with the same speed and resolution was recorded while phenylephrine (PE) (10 µM) was applied for the first 5 minutes and washed out using aCSF for the next 10 minutes.

### Image analysis

Image analysis was done as previously described by ^10^, using ImageJ and MATLAB R2020b (MathWork). For *in vivo* data analysis, neuronal somata regions of interest (ROIs) were hand-selected using ImageJ based on the RCaMP1.07 fluorescence. ROIs within astrocytes processes (*in vivo* and *in vitro* image analysis) or neuronal dendrites (*in vivo* image analysis) were automatically identified using an activity-based algorithm from a custom-designed image processing toolbox for MATLAB (Cellular and Hemodynamic Image Processing Suite (CHIPS) ^46^). Active pixels were defined based on two criteria relative to a sliding temporal boxcar of 5 s across the movie: 1) amplitude changes-active pixels exceeded 7 times the standard deviation (*in vivo* images, Lck-GCaMP6f AND RCaMP1.07) or 5 times the standard deviation (*in vitro* images, Lck-GCaMP6f) of the mean pixel intensity in this temporal boxcar and 2) timing-active pixels had a peak rise time within 0.07-1 s (*in vivo*, RCaMP1.07), 0.1-1 s (*in vivo* Lck-GCaMP6f) or 0.1-8 s (*in vitro*, Lck-GCaMP6f) compared to temporal boxcar. Active pixels were grouped within space (spatial radius of 4 µm) and time (0.2 s for RCaMP1.07 and 0.5 s Lck-GCaMP). The 3D mask of active pixels was summed along the temporal dimension, normalized, and thresholded (q = 0.2) to make a 2D activity ROI mask. Raw image data from pixels within each 2D ROI were statistically compared to pixels surrounding the ROI (p value < 0.05 by one-way ANOVA) to exclude false positives. Neuronal activity masks created by the algorithm and manually selected ROI mask were compared and the overlapping areas were excluded from the activity mask making sure each ROI was unique. For each ROI, a signal vector (dF/F) was calculated relative to the baseline fluorescence in the first 5 s of the trial and signal events were detected using the findpeaks function in MATLAB. For each event, different features such as amplitude and peak onset latency were measured and finally exported as .csv files for statistical analysis. The peak onset latency was calculated from the smoothed signal trace (5 frame moving average) as the first time point when the signal went over the threshold of 2.5 times the standard deviation of the baseline after the start of stimulation. We categorized astrocytes Ca^2+^ events as fast and delayed based on the median onset latency of their respective neurons during stimulation (control; 1.71s and Grin1 KD 1.78s). For all analysis, activities within the stimulation window (0 < onset latency < 8 s) were compared to the equivalent in trials with no stimulation except for fraction of active pixels and repeated response score which compared the events during the whole trial. For neurons, the amplitude distribution was described by a log normal distribution (R^2^ > 0.95). We categorized neurons based on this stable distribution using the 98^th^ and 88^th^ percentile of control neurons amplitude: High responding neurons (amplitude ≥ 3.47 dF/F), Mid responding neurons (1.92 ≤ amplitude <3.47 dF/F), and low responding neurons (amplitude < 1.92 dF/F). We also used a seed-based correlation analysis to correlate the signal vector (dF/F) for each ROI with the vectors from all other ROIs in the same field of view and examined the mean Pearson’s correlation coefficient across each trial within stimulation window (0-8s).

### Behavior tasks

The whisker touch-based novel texture recognition test and novel object recognition task (NORT) were performed to assess whisker discrimination or recognition memory ^4,59^ Both tasks included three general phases: habituation, learning, and testing. During the habituation phase, animals were introduced to an empty arena (40cm^3^) for 10 minutes a day for two consecutive days. The animal’s performance inside the arena was monitored and animals with stress signs (e.g., no exploration or moving around) were excluded. For the novel texture recognition test, following the last session of habituation, animals were lightly anesthetized using isoflurane (4% induction, 1-1.5% maintenance) and all the whiskers on the whisker plate ipsilateral to the chronic window were trimmed back to the face using fine-tipped scissors. On the contralateral side of the nose, all whiskers except for those corresponding to the virus injection area were also trimmed. During the learning phase of the novel texture recognition test, animals were introduced to the arena with two stands (4 cm × 15 cm) covered with the same grade of sandpaper, e.g. 150 grit, placed in the middle of the arena with an equal distance from each other and walls. Animals were allowed to explore for 5 minutes. After the learning phase, animals were transferred to a resting cage for 5 minutes. During this time, the stands with sandpaper were removed from the arena and two new stands were replaced: one familiar sandpaper (exactly same as the one in learning phase) and a novel sandpaper (e.g. 220 grit). During the testing phase, animals were allowed to explore the arena with one familiar and one novel texture for 3 minutes. The animal’s performance was recorded using a camera. To minimize the impact of olfactory cues, three copies of each stand were used. Also, the arena and the objects were cleaned with 70% ethanol between the learning and testing phase and between animals. The novel object recognition test was performed in the same manner, but without trimming the whiskers in advance and utilizing distinct objects, such as two 3D-printed dark blue bulldog shapes for the learning phase and a bulldog with an orange mouse shape for the testing phase.

### Behavior task analysis

Videos were analysed either manually or using a deep learning software, DeepLabCut (DLC ^90^), which allows for pose estimation of user-defined body parts using deep learning. DLC was first trained with 10 short clips (30 seconds) of the learning phase. The trained network was then used to estimate the position of animal’s nose as well as the objects in each frame of the video. Finally, a .csv file was created including the position of marked targets across videos. In either case, the amount of time animals spent around each object in each phase was measured. Investigation time was defined as the total time the animal was facing towards the sandpaper/object with their nose being at 2 cm or closer to the object. Climbing over the objects or stands was not considered as investigation. Any animal that did not explore any objects in the learning phase, explored only one object in the testing phase, or had a total investigation time of less than 2 s in the learning or testing phase was excluded from the analysis. The discrimination index was defined as the difference between time spent exploring novel texture vs. familiar texture divided by total exploration time (Novel-familiar exploration time/total exploration time).

### Fluorescence-Activated Cell Sorting (FACS)

Animals were anesthetised by intraperitoneal injection of pentobarbital (150 mg/Kg) and were transcardially perfused with Mg^2+^- and Ca^2+^-containing Hanks Buffered Salt Solution (HBSS^+Ca+Mg^) (NaCl, 140 mM; KCl, 5 mM; MgCl_2_*6H2O, 5 mM; MgSO4-7H2O, 0.4 mM; CaCl_2_, 1 mM; Na2HPO4-2H2O, 0.3 mM; NaHCO_3_, 4 mM; KH2PO4, 0.4 mM; D-Glucose, 6 mM). The injection area under the cranial window was cut into small pieces and was incubated with Dispase II (final concentration of 0.6U/ml; Millipore-Sigma; D4693) incubated at 37°C for 1 hour with gentle shaking (120 rpm). Then, the tissue was gently dissociated by passing through a 1ml pipette tip for 7 times, followed by a 40µm pipette tip cell strainer (SP Bel-Art Flowmi). Cells were then spun at 400xg at 4°C for 5 minutes. Cells were washed twice with ice-cold HBSS^+Ca+Mg^ at 400xg for 5 minutes removing supernatant between each wash and keeping the pellet untouched. Cells were gently resuspended in ice-cold HBSS^+Ca+Mg^ containing DNaseI (5U; Fisher Scientific; RQ1) and placed on ice until FACS. Single cell sorting using a FACS machine (BD FACSAriaIII) was conducted by technicians at the Flow Cytometry core facility at the University of Manitoba. Astrocyte Lck-GCaMP fluorescence (i.e. green GFP fluorescence) was used for sorting. FACSDiva (Version 6.1.3) software was used for analysis of the recorded FACS events. For each signal, the side scatter was plotted against the fluorescence intensity.

### Quantification of Grin1 expression using qRT-PCR

Upon sorting, cells were lysed using the RNA Purification kit (ThermoFisher) according to manufacturer’s instructions and were stored at −80°C until RNA extraction. RNA was extracted from sorted GCaMP6f-expressing astrocytes using the Purelink RNA Micro Kit (ThermoFisher) according to manufacturer’s instructions. RNA amplification was done using MessageAmp II aRNA Amplification Kit (ThermoFisher) according to manufacturer’s instructions except for the aRNA purification where we used Trizol to increase the recovery rate. Reverse transcription of aRNA was done according to SuperScript IV VILO (ThermoFisher). We used Qiagen PCR Purification kit for cDNA purification. Real-time PCR samples were prepared by using the purified cDNA (5uL; 2ng/uL), PowerUp SYBR MasterMix (Life Technologies; A25742) (10 µl), and primers (1µl F and 1 µl R) which were all designed using NCBI primer blast software. PCR was done in triplicate on the QuantStudio 6 Flex with the following program: 2 minutes at 50°C, 2 minutes at 95°C, 45X 5 seconds at 95°C, and 30 seconds at 60°C. Analysis was done in QuantStudio 6 Flex software. All samples were gender and age matched (age of injection and days post injection, see table 1). Using Normfinder and Bestkeeper softwares, 3 of the housekeeping genes found to be the most stable were used in the final analysis including Actb, Atp5pb and Hprt. Relative expression of Grin1 gene in Grin1 KD astrocytes was calculated based on previously generated standard curves (8-point dilution series), normalized to the expression of the 3 housekeeping genes, and finally compared to the average non-silencing relative gene expression. Primers sequences are as follows:

MusHPRT F667: 5’-ACAGGCCAGACTTTGTTGGA-3’; MusHPRT R765: 5’-CACAAACGTGATTCAAATCCCTGA-3’; MusACTB F902: 5’-TCCTTCTTGGGTATGGAATCCTG-3’; MusACTB R987: 5’-AGGTCTTTACGGATGTCAACG-3’; Atp5pb F; 5’-GTCCAGGGGTATTACAGGCAA-3’; Atp5pb R: 5’-TCAGGAATCAGCCCAAGACG-3’; Ywhaz F: 5’-ATCCCCAATGCTTCGCAACC-3’; Ywhaz R: 5’-ACTGGTCCACAATTCCTTTCTTG-3’; MusGrin1 1237F: 5’-CAACATCTGGAAGACAGGACC-3’, MusGrin1 1308R: 5’-CCAGTCACTCCATCTGCATAC-3’; Tubb3 F: 5’-ACCATGGACAGTGTTCGGTC-3’; Tubb3 R: 5’-AGCACCACTCTGACCAAAGATA-3’; S100b F: 5’-CTTCCTGCTCCTTGATTTCCTCCA-3’; S100b R: 5’-CGAGAGGGTGACAAGCACAAG-3’.

### Immunohistochemistry

Animals were anesthetised by intraperitoneal injection of pentobarbital (150 mg/Kg) and were transcardially perfused with 20 mL oxygenated aCSF followed by 60 ml 2% paraformaldehyde (PFA; Millipore-Sigma; P6148) (in 2X Phosphate-buffered saline (PBS), pH adjusted to 7.2-7.4). Hemispheres were then post-fixed in 4% PFA for 3 hours, washed with ice-cold 1X PBS, and cryoprotected with 30% sucrose (in 1X PBS) in 4°C overnight. Tissue mounted on Optimal cutting temperature compound (OCT compound; Tissue Tek) was cut on a cryostat (Leica) to obtain 40 µm sections. Sections were washed twice with TBST (Tris-buffered saline+ polysorbate/Tween 20) for 10 minutes while incubated on a rocking platform at room temperature. Sections were then incubated in blocking solution including TBS + 0.3% Triton X-100 (0.3% TBST) + 5% Donkey Normal Serum (DNS) + IgG Fab blocking reagent (90uL/ml; Jackson Immunoresearch) for 1 hour. Subsequently, sections were washed 3 times with TBST for 5 minutes. Sections were incubated in 0.3% TBST + 1% NDS + primary antibodies overnight at 4°C on a rocking platform. Following incubation with primary antibodies, sections were washed with 0.05% TBST 3 times for 10 minutes and were incubated with secondary antibodies and 0.05% TBST for 1 hour at room temperature. Following incubation, sections were washed twice with 0.05% TBST and once with TBS for 10 minutes. Sections were then washed in 0.1X PBS and mounted on glass slides with a coverslip. Images of sections were acquired on a confocal microscope (Zeiss LSM 810). Primary antibodies include Chicken-anti-GFP (1:1000, Aves), Rabbit-anti-mouse/rat-GFAP (1:3000, Dako), and Mouse-anti-mouse/rat-NeuN (1;200, Millipore). Secondary antibodies include Goat-anti-Chicken-IgG-488 (1:1000, Invitrogen), Donkey-anti-Mouse-IgG-647 (1:1000, Invitrogen), and Donkey-anti-Rabbit-IgG-568 (1:1000, Invitrogen).

### Statistical analysis

All statistics for Ca^2+^ analysis was performed in R (Version 1.2.1335) using the lme4 package for linear mixed-effects models. For *in vivo* Ca^2+^ analysis, our fixed effects included animal type (control or Grin1 KD), stimulus condition (with or without stimulation), astrocyte type (fast or delayed), neuronal populations (High or mid or low responding neurons), and sex (male or female) as well as the interaction of these effects. As random effects, we had intercepts for individual animals, fields of view, ROIs, and trials. For *in vitro* Ca^2+^ analysis, fixed effects included type (control or Grin1 KD), stimulus condition (Drug or baseline) and the interaction between these two. Likelihood ratio tests comparing models with fixed effects against models without fixed effects were performed to determine the model with the best fit while accounting for the different degrees of freedom. All data were reported and plotted as uncorrected means and standard error of the means. P values for different parameter comparisons were obtained using the multcomp package with Tukey post hoc tests. Non-parametric Mann-Whitney-Wilcoxon tests were performed in R (Version 1.2.1335) for qPCR, immunostaining quantification and behavior tasks.

### Resource Availability

The code used for calcium analysis (CHIPS v1.1.0) is available as a MATLAB toolbox at (//github.com/EIN-lab/CHIPS/releases). AAVs for astrocyte Grin1 KD and control will be available upon publication of this manuscript at the University of Zurich Viral Vector Facility. Requests for further information, reagents, data and resources should be directed to and will be fulfilled by Dr. Jill Stobart upon publication of this manuscript.

## Acknowledgements

We would like to thank the staff in the Central Animal Care Services and Veterinary Services at the University of Manitoba for their diligent work and dedication to animal care and husbandry. This work was supported by a Discovery Grant from the Natural Sciences and Engineering Research Council of Canada (NSERC), the University Research Grants Program (University of Manitoba), and start-up funding from the University of Manitoba. The Stobart lab has also received support from the Canadian Institutes for Health Research, Research Manitoba, Brain Canada/Azrieli Foundation, and the Manitoba Medical Service Foundation. M.K. is supported by graduate studentships from the Rady Faculty of Health Sciences and Research Manitoba. T.S. was supported by a Mitacs Globalink Internship. D.E. was supported by an Undergraduate Research Award (University of Manitoba).

## Author Contributions

M.S. and J.S. designed the study. N.A., M.K., M.S., T.S., F.O., D.E., S.C-F. performed the experiments. N.A., M.K., M.S., and J.S. analyzed data. N.A., M.K. and J.S. wrote the paper.

## References

1. Mu, Y. et al. Glia accumulate evidence that actions are futile and suppress unsuccessful behavior. Cell 178, 27–43.e19 (2019).

2. Adamsky, A. et al. Astrocytic activation generates de novo neuronal potentiation and memory enhancement. Cell 174, 59–71.e14 (2018).

3. Yu, X. et al. Reducing astrocyte calcium signaling in vivo alters striatal microcircuits and causes repetitive behavior. Neuron 99, 1170–1187 (2018).

4. Kwak, H. et al. Astrocytes control sensory acuity via tonic inhibition in the thalamus. Neuron 108, 691–706.e10 (2020).

5. Nagai, J. et al. Specific and behaviorally consequential astrocyte Gq GPCR signaling attenuation in vivo with iβARK. Neuron 1–19 (2021). doi:10.1016/j.neuron.2021.05.023

6. Araque, A. et al. Gliotransmitters travel in time and space. Neuron 81, 728–739 (2014).

7. Bazargani, N. & Attwell, D. Astrocyte calcium signaling: the third wave. Nat. Neurosci. 19, 182–189 (2016).

8. Giaume, C., Koulakoff, A., Roux, L., Holcman, D. & Rouach, N. Astroglial networks: a step further in neuroglial and gliovascular interactions. Nat Rev Neurosci 11, 87–99 (2010).

9. Kastanenka, K. V. et al. A roadmap to integrate astrocytes into Systems Neuroscience. Glia 68, 5–26 (2020).

10. Stobart, J. L. et al. Cortical circuit activity evokes rapid astrocyte calcium signals on a similar timescale to neurons. Neuron 98, 726–735.e4 (2018).

11. Bindocci, E. et al. Three-dimensional Ca^2+^ imaging advances understanding of astrocyte biology. Science 356, eaai8185 (2017).

12. Srinivasan, R. et al. Ca^2+^ signaling in astrocytes from Ip3r2 −/− mice in brain slices and during startle responses in vivo. Nat. Neurosci. 18, 708–717 (2015).

13. Stobart, J. L. et al. Long-term in vivo calcium imaging of astrocytes reveals distinct cellular compartment responses to sensory stimulation. Cereb. Cortex 28, 184–198 (2018).

14. Shigetomi, E., Tong, X., Kwan, K. Y., Corey, D. P. & Khakh, B. S. TRPA1 channels regulate astrocyte resting calcium and inhibitory synapse efficacy through GAT-3. Nat. Neurosci. 15, 70–80 (2011).

15. Georgiou, L., Echeverría, A., Georgiou, A. & Kuhn, B. Ca+ activity maps of astrocytes tagged by axoastrocytic AAV transfer. Sci. Adv. 8, (2022).

16. Nizar, K. et al. *In vivo* stimulus-induced vasodilation occurs without IP_3_ receptor activation and may precede astrocytic calcium increase. J. Neurosci. 33, 8411–8422 (2013).

17. Tran, C. H. T., Peringod, G. & Gordon, G. R. Astrocytes integrate behavioral state and vascular signals during functional hyperemia. Neuron 100, 1–16 (2018).

18. Chen, N. et al. Nucleus basalis-enabled stimulus-specific plasticity in the visual cortex is mediated by astrocytes. Proc. Natl. Acad. Sci. U. S. A. 109, E2832–41 (2012).

19. Ding, F. et al. α1-adrenergic receptors mediate coordinated Ca^2+^ signaling of cortical astrocytes in awake, behaving mice. Cell Calcium 54, 387–394 (2013).

20. Paukert, M. et al. Norepinephrine controls astroglial responsiveness to local circuit activity. Neuron 82, 1263–1270 (2014).

21. Agarwal, A. et al. Transient opening of the mitochondrial permeability transition pore induces microdomain calcium transients in astrocyte processes. Neuron 93, 587–605.e7 (2017).

22. Petravicz, J., Fiacco, T. a & McCarthy, K. D. Loss of IP3 receptor-dependent Ca^2+^ increases in hippocampal astrocytes does not affect baseline CA1 pyramidal neuron synaptic activity. J. Neurosci. 28, 4967–4973 (2008).

23. Takata, N. et al. Astrocyte calcium signaling transforms cholinergic modulation to cortical plasticity in vivo. J. Neurosci. 31, 18155–18165 (2011).

24. Rungta, R. L. et al. Ca^2+^ transients in astrocyte fine processes occur via Ca^2+^ influx in the adult mouse hippocampus. Glia 64, 2093–2103 (2016).

25. Shigetomi, E., Jackson-Weaver, O., Huckstepp, R. T., O’Dell, T. J. & Khakh, B. S. TRPA1 channels are regulators of astrocyte basal calcium levels and long-term potentiation via constitutive d-serine release. J. Neurosci. 33, 10143–10153 (2013).

26. Ziemens, D., Oschmann, F., Gerkau, N. J. & Rose, C. R. Heterogeneity of activity-induced sodium transients between astrocytes of the mouse hippocampus and neocortex: mechanisms and consequences. J. Neurosci. 39, 2620–2634 (2019).

27. Letellier, M. et al. Astrocytes regulate heterogeneity of presynaptic strengths in hippocampal networks. Proc. Natl. Acad. Sci. U. S. A. 113, E2685–E2694 (2016).

28. Kondoh, T., Nishizaki, T., Aihara, H. & Tamaki, N. NMDA-responsible, APV-insensitive receptor in cultured human astrocytes. Life Sci. 68, 1761–1767 (2001).

29. Skowronska, K., Obara-Michlewska, M., Zielinska, M. & Albrecht, J. NMDA Receptors in Astrocytes : In search for roles in neurotransmission and astrocytic homeostasis. Int. J. Mol. Sci. 20, (2019).

30. Jimenez-Blasco, D., Santofimia-Castanõ, P., Gonzalez, A., Almeida, A. & Bolanõs, J. P. Astrocyte NMDA receptors’ activity sustains neuronal survival through a Cdk5-Nrf2 pathway. Cell Death Differ. 22, 1877–1889 (2015).

31. Serrano, A., Robitaille, R. & Lacaille, J. C. Differential NMDA-dependent activation of glial cells in mouse hippocampus. Glia 56, 1648–1663 (2008).

32. Montes de Oca Balderas, P. & Gonzalez Hernandez, J. R. NMDA receptors in astroglia: Chronology, controversies, and contradictions from a complex molecule. in Astrocyte-Physiology and Pathology 51–89 (InTech Open, 2018). doi:http://dx.doi.org/10.5772/57353

33. Schipke, C. G. et al. Astrocytes of the mouse neocortex express functional N-methyl-D-aspartate receptors. FASEB J. 15, 1270–1272 (2001).

34. Verkhratsky, A. & Chvátal, A. NMDA Receptors in Astrocytes. Neurochem. Res. 45, 122–133 (2020).

35. Lalo, U., Pankratov, Y., Parpura, V. & Verkhratsky, A. Ionotropic receptors in neuronal-astroglial signalling: What is the role of ‘excitable’ molecules in non-excitable cells. Biochim. Biophys. Acta 1813, 992–1002 (2011).

36. Lalo, U., Pankratov, Y., Kirchhoff, F., North, R. A. & Verkhratsky, A. NMDA receptors mediate neuron-to-glia signaling in mouse cortical astrocytes. J. Neurosci. 26, 2673–2683 (2006).

37. Palygin, O., Lalo, U., Verkhratsky, A. & Pankratov, Y. Ionotropic NMDA and P2X1/5 receptors mediate synaptically induced Ca^2+^ signalling in cortical astrocytes. Cell Calcium 48, 225–231 (2010).

38. Palygin, O., Lalo, U. & Pankratov, Y. Distinct pharmacological and functional properties of NMDA receptors in mouse cortical astrocytes. Br. J. Pharmacol. 163, 1755–1766 (2011).

39. Cahoy, J. D. et al. A transcriptome database for astrocytes, neurons, and oligodendrocytes: A new resource for understanding brain development and function. J. Neurosci. 28, 264–278 (2008).

40. Lalo, U., Palygin, O., North, R. A., Verkhratsky, A. & Pankratov, Y. Age-dependent remodelling of ionotropic signalling in cortical astroglia. Aging Cell 10, 392–402 (2011).

41. Haustein, M. D. et al. Conditions and constraints for astrocyte calcium signaling in the hippocampal mossy fiber pathway. Neuron 82, 413–429 (2014).

42. Otsu, Y. et al. Calcium dynamics in astrocyte processes during neurovascular coupling. Nat. Neurosci. 18, 210–218 (2015).

43. Lind, B. L. et al. Fast Ca^2+^ responses in astrocyte end-feet and neurovascular coupling in mice. Glia 66, 348–358 (2018).

44. Institoris, A. et al. Astrocytes amplify neurovascular coupling to sustained activation of neocortex in awake mice. (2022). doi:10.1038/s41467-022-35383-2

45. Huntington, T. E. & Srinivasan, R. Astrocytic mitochondria in adult mouse brain slices show spontaneous calcium influx events with unique properties. Cell Calcium 96, 102383 (2021).

46. Barrett, M. J. P., Ferrari, K. D., Stobart, J. L., Holub, M. & Weber, B. CHIPS: an extensible toolbox for cellular and hemodynamic two-photon image analysis. Neuroinformatics 16, 145–147 (2018).

47. Glück, C. et al. Distinct signatures of calcium activity in brain mural cells. Elife 10, 1–27 (2021).

48. Margolis, D. J. et al. Reorganization of cortical population activity imaged throughout long-term sensory deprivation. Nat. Neurosci. 15, 1539–1546 (2012).

49. Mayrhofer, J. M., Haiss, F., Helmchen, F. & Weber, B. Sparse, reliable, and long-term stable representation of periodic whisker deflections in the mouse barrel cortex. Neuroimage 115, 52–63 (2015).

50. Lefort, S., Tomm, C., Floyd Sarria, J. C. & Petersen, C. C. H. The excitatory neuronal network of the C2 barrel column in mouse primary somatosensory cortex. Neuron 61, 301–316 (2009).

51. Petersen, C. C. H. & Crochet, S. Synaptic computation and sensory processing in neocortical layer 2/3. Neuron 78, 28–48 (2013).

52. Sakata, S. & Harris, K. D. Laminar structure of spontaneous and sensory-evoked population activity in auditory cortex. Neuron 64, 404–418 (2009).

53. O’Connor, D. H., Peron, S. P., Huber, D. & Svoboda, K. Neural activity in barrel cortex underlying vibrissa-based object localization in mice. Neuron 67, 1048–1061 (2010).

54. Kerr, J. N. D. et al. Spatial organization of neuronal population responses in Layer 2/3 of rat barrel cortex. J. Neurosci. 27, 13316–13328 (2007).

55. Zhao, J., Wang, D. & Wang, J.-H. Barrel cortical neurons and astrocytes coordinately respond to an increased whisker stimulus frequency. Mol. Brain 5, 12 (2012).

56. Nizar, K. et al. In vivo stimulus-induced vasodilation occurs without IP3 receptor activation and may precede astrocytic calcium increase. J. Neurosci. 33, 8411–8422 (2013).

57. Wang, X. et al. Astrocytic Ca^2+^ signaling evoked by sensory stimulation *in vivo*. Nat. Neurosci. 9, 816–823 (2006).

58. Lind, B. L., Brazhe, A. R., Jessen, S. B., Tan, F. C. C. & Lauritzen, M. J. Rapid stimulus-evoked astrocyte Ca^2+^ elevations and hemodynamic responses in mouse somatosensory cortex in vivo. Proc. Natl. Acad. Sci. U. S. A. 110, E4678–E4687 (2013).

59. Wu, H. P. P., Ioffe, J. C., Iverson, M. M., Boon, J. M. & Dyck, R. H. Novel, whisker-dependent texture discrimination task for mice. Behav. Brain Res. 237, 238–242 (2013).

60. Ahmadpour, N., Kantroo, M. & Stobart, J. L. Extracellular calcium influx pathways in astrocyte calcium microdomain physiology. (2021).

61. Hogan-Cann, A. D. & Anderson, C. M. Physiological roles of non-neuronal NMDA receptors. Trends Pharmacol. Sci. 37, 750–767 (2016).

62. Kol, A. et al. Astrocytes contribute to remote memory formation by modulating hippocampal–cortical communication during learning. Nat. Neurosci. 23, 1229–1239 (2020).

63. Lines, J., Martin, E. D., Kofuji, P., Aguilar, J. & Araque, A. Astrocytes modulate sensory-evoked neuronal network activity. Nat. Commun. 11, 3689 (2020).

64. Benz, B., Grima, G. & Do, K. Q. Glutamate-induced homocysteic acid release from astrocytes: possible implication in glia-neuron signaling. Neuroscience 124, 377–386 (2004).

65. Gérard, F. & Hansson, E. Inflammatory activation enhances NMDA-triggered Ca^2+^ signalling and IL-1β secretion in primary cultures of rat astrocytes. Brain Res. (2012). doi:10.1016/j.brainres.2012.07.032

66. Montes de Oca Balderas, P. & Aguilera, P. A metabotropic-Like flux-Independent NMDA receptor regulates ca^2+^ exit from endoplasmic reticulum and mitochondrial membrane potential in cultured astrocytes. PLoS One 10, 1–22 (2015).

67. Zhang, Q., Hu, B., Sun, S. & Tong, E. Induction of increased intracellular calcium in astrocytes by glutamate through activating NMDA and AMPA receptors. J. Huazhong Univ. Sci. Technolog. Med. Sci. 23, 254–257 (2003).

68. Hu, B., Sun, S. G. & Tong, E. T. NMDA and AMPA receptors mediate intracellular calcium increase in rat cortical astrocytes. Acta Pharmacol. Sin. 25, 714–720 (2004).

69. Mehina, E. M. F., Murphy-Royal, C. & Gordon, G. R. Steady-state free Ca^2+^ in astrocytes is decreased by experience and impacts arteriole tone. J. Neurosci. 37, 8150–8165 (2017).

70. Avermann, M., Tomm, C., Mateo, C., Gerstner, W. & Petersen, C. C. H. Microcircuits of excitatory and inhibitory neurons in layer 2/3 of mouse barrel cortex. J. Neurophysiol. 107, 3116–3134 (2012).

71. Holmgren, C., Harkany, T., Svennenfors, B. & Zilberter, Y. Pyramidal cell communication within local networks in layer 2/3 of rat neocortex. J. Physiol. 551, 139 (2003).

72. Petersen, C. C. H. Sensorimotor processing in the rodent barrel cortex. Nat. Rev. Neurosci. 20, 533–546 (2019).

73. Adesnik, H., Bruns, W., Taniguchi, H., Huang, Z. J. & Scanziani, M. A neural circuit for spatial summation in visual cortex. Nature 490, 226 (2012).

74. Gentet, L. J. et al. Unique functional properties of somatostatin-expressing GABAergic neurons in mouse barrel cortex. Nat. Neurosci. 2012 154 15, 607–612 (2012).

75. Gentet, L. J. Functional diversity of supragranular GABAergic neurons in the barrel cortex. Front. Neural Circuits 6, 1–13 (2012).

76. Karnani, M. M. et al. Opening holes in the blanket of inhibition: Localized lateral disinhibition by VIP interneurons. J. Neurosci. 36, 3471 (2016).

77. Crochet, S. & Petersen, C. C. H. Correlating whisker behavior with membrane potential in barrel cortex of awake mice. Nat. Neurosci. 9, 608–610 (2006).

78. Poulet, J. F. A. & Petersen, C. C. H. Internal brain state regulates membrane potential synchrony in barrel cortex of behaving mice. Nat. 2008 4547206 454, 881–885 (2008).

79. Kyriakatos, A. et al. Voltage-sensitive dye imaging of mouse neocortex during a whisker detection task. Neurophotonics 4, 031204 (2017).

80. Brecht, M., Schneider, M. & Manns, I. D. Silent neurons in sensorimotor cortices: Implications for cortical Plasticity. in *In* F. F. Ebner (Ed.), Neural plasticity in adult somatic sensory-motor systems 1–19 (2005).

81. Butcher, J. B. et al. A requirement for astrocyte IP 3 R2 signaling for whisker depression and homeostatic upregulation in the mouse barrel cortex. Front. Cell. Neurosci. 16, (2022).

82. Chipman, P. H. et al. Astrocyte GluN2C NMDA receptors control basal synaptic strengths of hippocampal CA1 pyramidal neurons in the stratum radiatum. Elife 10, 1–32 (2021).

83. Chung, S., Li, X. & Nelson, S. B. Short-term depression at thalamocortical synapses contributes to rapid adaptation of cortical sensory responses in vivo. Neuron 34, 437–446 (2002).

84. Khatri, V., Hartings, J. A. & Simons, D. J. Adaptation in thalamic barreloid and cortical barrel neurons to periodic whisker deflections varying in frequency and velocity. J. Neurophysiol. 92, 3244–3254 (2004).

85. Adibi, M., McDonald, J. S., Clifford, C. W. G. & Arabzadeh, E. Adaptation improves neural coding efficiency despite increasing correlations in variability. J. Neurosci. 33, 2108–2120 (2013).

86. Musall, S. et al. Tactile frequency discrimination is enhanced by circumventing neocortical adaptation. Nat. Neurosci. 17, 1567–1573 (2014).

87. Tannan, V., Simons, S., Dennis, R. G. & Tommerdahl, M. Effects of adaptation on the capacity to differentiate simultaneously delivered dual-site vibrotactile stimuli. Brain Res. 1186, 164–170 (2007).

88. Lalo, U. et al. Exocytosis of ATP from astrocytes modulates phasic and tonic inhibition in the neocortex. PLoS Biol. 12, e1001747 (2014).

89. Fellmann, C. et al. An optimized microRNA backbone for effective single-copy RNAi. Cell Rep. 5, 1704–1713 (2013).

90. Mathis, A. et al. DeepLabCut: markerless pose estimation of user-defined body parts with deep learning. Nat. Neurosci. 21, 1281–1289 (2018).

