## Supplementary Figures for "Cortical astrocyte N-Methyl-D-Aspartate receptors influence whisker barrel activity and sensory discrimination"

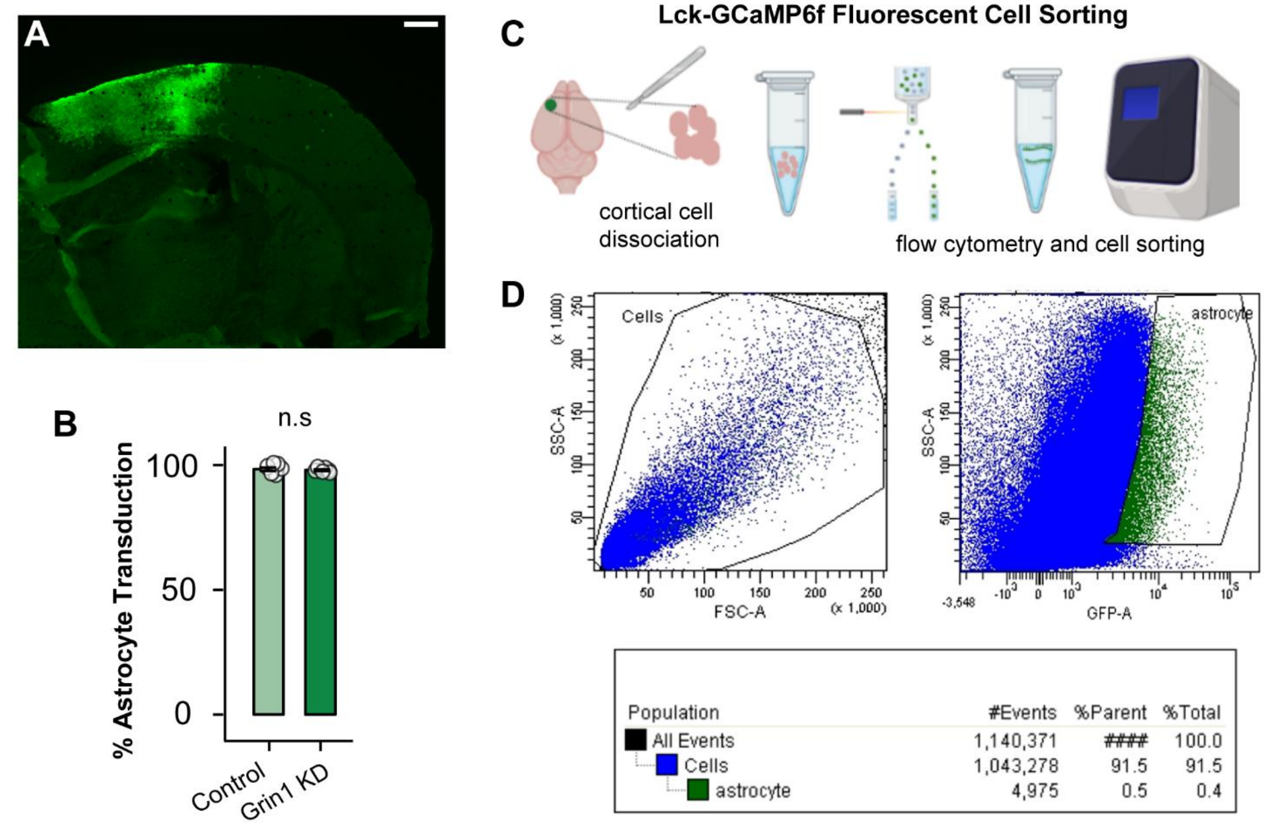


**Supplementary Figure 1. Astrocyte viral transduction and FACS.** A) The astrocyte viral constructs (green, stained with anti-GFP) labelled a good section of cortex. Scale bar= 500 µm. B) The viruses transduced more than 95% of GFAP positive astrocytes in the injection area. C) Schematic for cortical cell dissociation and fluorescence-associated cell sorting (FACS) using Lck-GCaMP6f fluorescence. D) Example dot plots from the flow cytometer with non-fluorescent cells (blue) and the collected GFP population (green). Astrocytes were selected based on GCaMP (GFP) fluorescence and large granularity (SSC-A). Statistics were calculated by Mann-Whitney-Wilcoxon test.

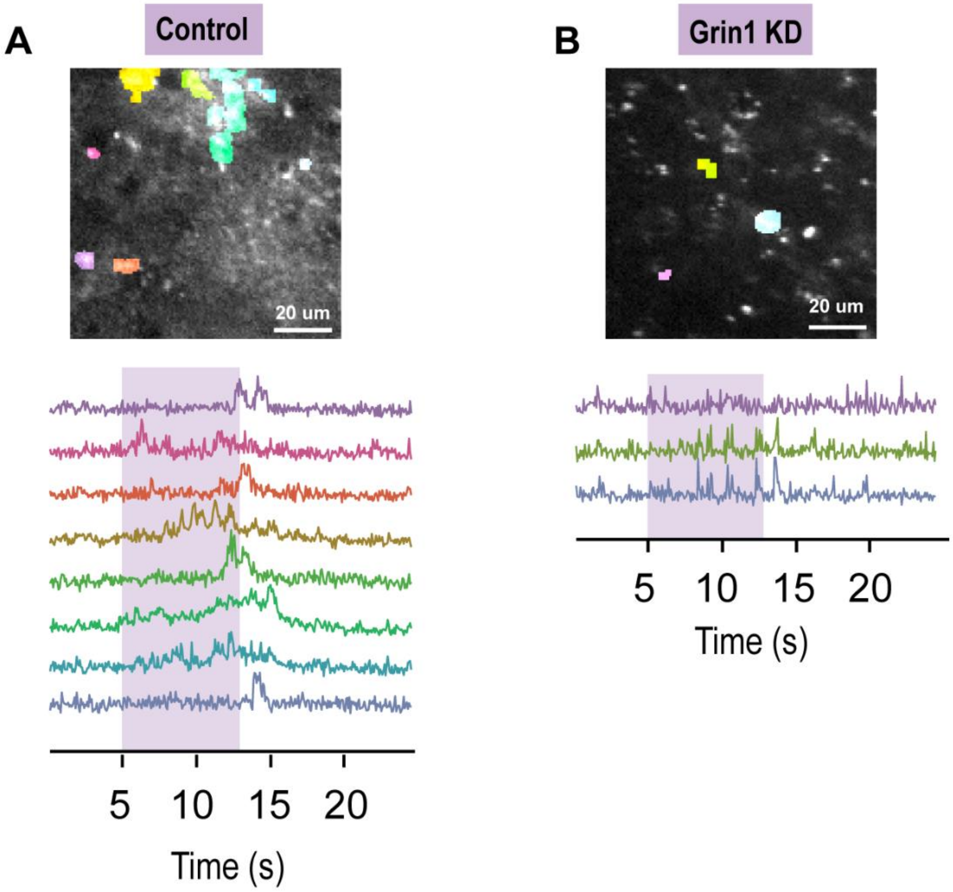


**Supplementary Figure 2. Astrocyte stimulus-evoked $Ca^{2+}$ responses.** Example traces from ROIs evoked during whisker stimulation (purple box) in control (A) and Grin 1 KD (B).

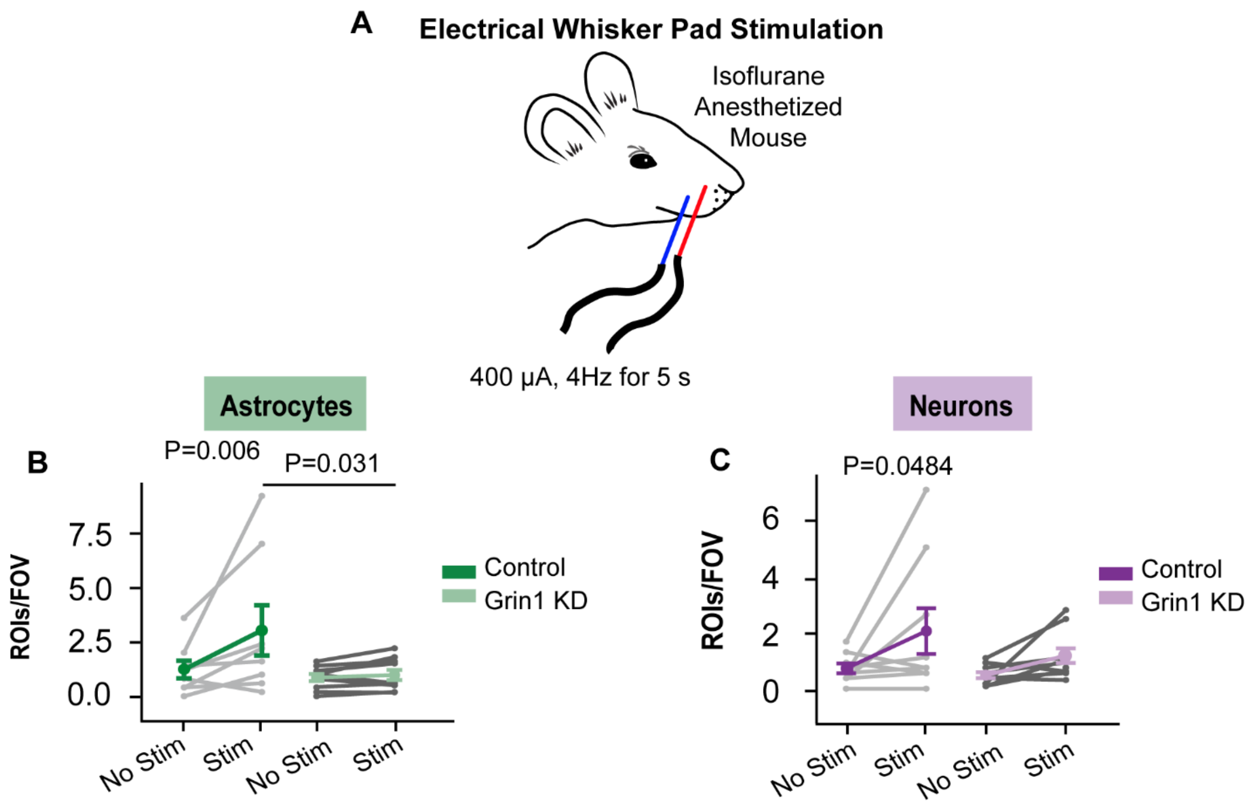


**Supplementary Figure 3. Astrocyte and neuron responses to brief electrical whisker pad stimulation in anesthetized mice.** A) Experimental schematic. B) Number of astrocyte microdomain ROIs per FOV evoked by electrical stimulation. C) Number of neuron ROIs per FOV evoked by electrical stimulation. n= 8 control and 10 Grin1 KD mice. Statistics were calculated using linear mixed model and Tukey post hoc tests.

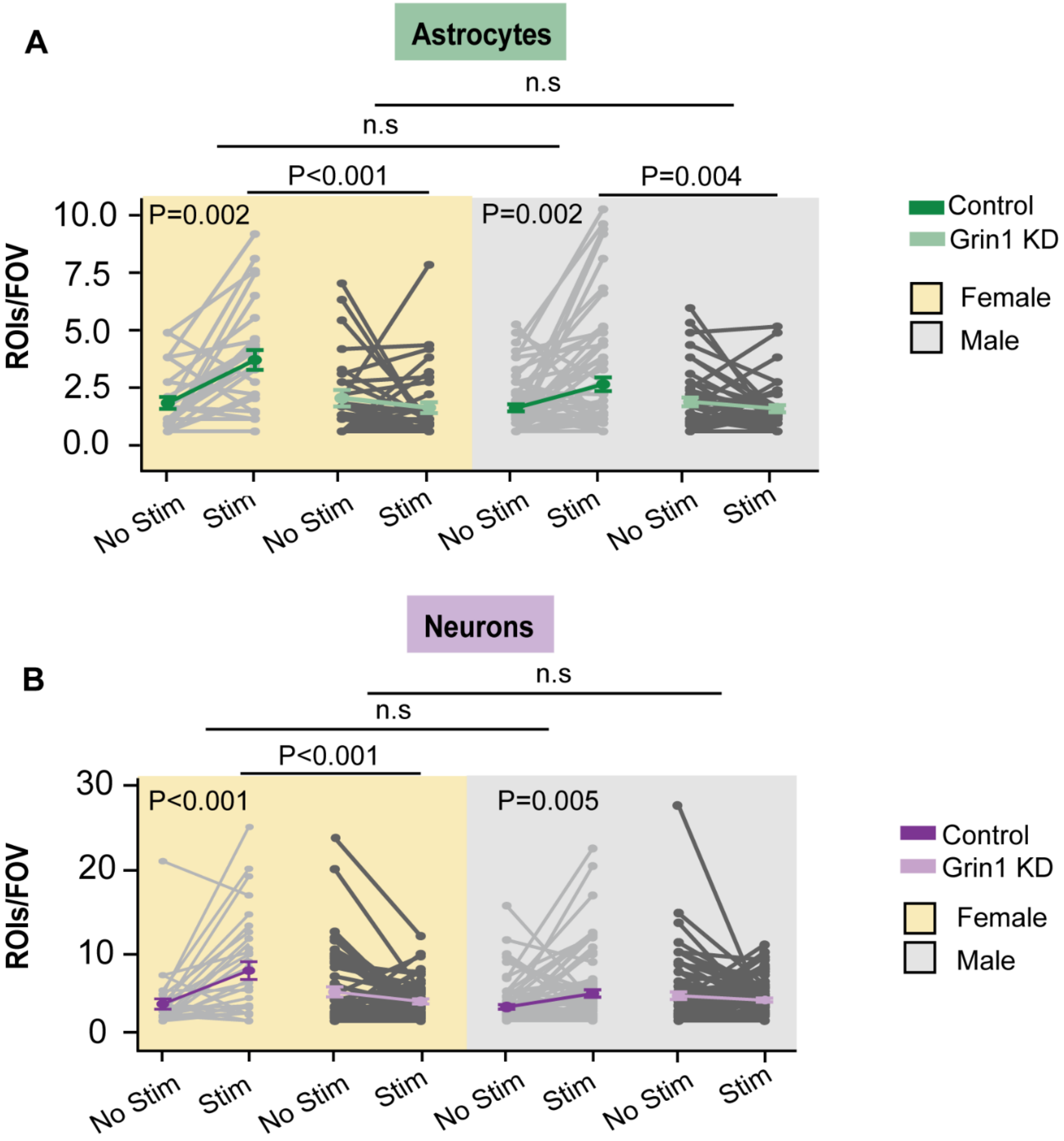


**Supplementary Figure 4. Sex comparisons for astrocyte and neurons $Ca^{2+}$ responses.** A) Mean number of ROIs/FOV in Lck-GCaMP6f astrocytes from male and female mice. B) Mean number of ROIs/FOV in RCaMP neurons from male and female mice. No sex differences were detected in any case. Control: n=103 FOV, 4 female and 4 male, Grin1 KD: n=97 FOV, 6 female mice, 5 male mice. Statistics were calculated using linear mixed model and Tukey post hoc tests.
